# Potential beneficial effects of PD-1/PD-L1 blockade in Alzheimer’s disease: A systematic review and meta-analysis of preclinical and clinical studies

**DOI:** 10.1101/2025.06.18.660247

**Authors:** Jiyoung Yoon, Heonyoung Ha, Hyun Woo Lee, Seungyeon Kim, Yun Mi Yu, Heejung Chun

## Abstract

Programmed cell death-1 (PD-1) and its ligand (PD-L1) play key roles in cancer immune evasion and in modulating neuroinflammation. This systematic review and meta-analysis investigated the effect of PD-1/PD-L1 blockade on pathology and cognitive function in Alzheimer’s disease (AD) in preclinical and clinical studies. Relevant studies were systematically identified using the MEDLINE, Embase, CENTRAL, and Web of Science databases from their inception until April 10, 2024. In total, 33 studies were included in this meta-analysis, conducted using R software. Preclinical studies revealed that blockade of PD-1 signaling reduces amyloid-beta plaque burden, tau phosphorylation, and astrocyte reactivity in AD mouse models, accompanied by improvements in cognitive function in behavioral tests. Furthermore, clinical studies demonstrated the beneficial effect of PD-1 signaling inhibitors on cognitive function in cancer patients. This study highlights the necessity for additional research to clarify the exact mechanisms by which PD-1/PD-L1 inhibition impacts AD pathology and cognitive function, opening avenues for potential therapeutic strategies targeting this pathway in AD.

## Introduction

Alzheimer’s disease (AD) is the primary cause of dementia and lacks effective treatment options at present. The main hallmarks of AD are amyloid-beta (Aβ) deposits, reactive astrocytes, and aggregation of phosphorylated tau, resulting in neurofibrillary tangles [1, 2]. Aβ deposits induce reactive astrocytes, which link early-stage Aβ aggregation [3–5], with late-stage tau pathology in AD [6–9]. Reactive astrocytes also trigger the secretion of pro-inflammatory factors, leading to a neuroinflammatory milieu [5–7]. Under neuroinflammatory conditions, resident-brain immune cells, such as microglia, are activated. The structural stability of the blood-brain barrier (BBB) is also impaired, with a subsequent increase in the infiltration of peripheral immune cells into the brain [10, 11]. It has been reported that the infiltrated regulatory T (Treg) cells or monocyte-derived macrophages (MDMs) significantly attenuate of neurodegeneration and cognitive decline in AD mouse models [12–14]. However, the mechanisms underlying this alleviation remain unclear.

The ability of brain-resident or infiltrating immune cells to alleviate AD pathology may be potentially influenced by the signaling of the inhibitory receptor programmed cell death protein-1 (PD-1) and its ligand, programmed cell death-ligand 1 (PD-L1). This signaling has been extensively studied for its crucial role in enabling tumors to evade immune surveillance. PD-1 is found on immune cells, including activated T cells, and tumor-associated macrophages. It interacts with PD-L1 which is expressed in various cells, including those in tumors. PD-1/PD-L1 axis suppresses T cell activity, enabling cancer cells to exploit the immune system. Therefore, the interruption of this receptor/ligand pathway by inhibitors enhances anti-tumor immune response [15, 16]. In the central nervous system (CNS), PD-1/PD-L1 signaling has been associated with multifaceted function. Under physiological conditions, PD-1 is primarily expressed in neurons, notably in the hippocampal CA1 and CA3 [17] and dorsal root ganglion sensory [18]. The PD-1 receptor expressed in the hippocampal neurons plays a role in regulating neuronal activity and synaptic plasticity [19]. In contrast, under pathological conditions, PD-1 expression is elevated in infiltrating immune cells or microglia, and PD-L1 expression is escalated in reactive astrocytes [20, 21]. Although PD-1/PD-L1 axis is deemed to regulate the immune system in the brain [22, 23], the function of PD-1/PD-L1 axis in the CNS is yet to be fully elucidated.

In clinical settings, PD-1/PD-L1 inhibitors are increasingly utilized to manage various cancer types, with expanding indications owing to their anticancer effects [24, 25]. Numerous clinical studies are investigating the anticancer effects of such inhibitors in patients with oncological disorders. Additionally, there is a continuous flow of research evaluating the safety profiles of these inhibitors, consistently addressing cognitive impairment, which is a potential side effect of most chemotherapies. Recent clinical studies have shown that fewer adverse cognitive effects were caused by PD-1/PD-L1 inhibitors compared with conventional chemotherapy [26, 27]. However, some studies have failed to detect a notable difference in the cognitive impact between these inhibitors and conventional chemotherapy [28–30]. Given the inconsistencies across preclinical and clinical studies and the limited empirical research on the correlation between the blockade of PD-1 signaling and cognitive function, a systematic review and meta-analysis of relevant studies is required to provide comprehensive examination of the effects of PD-1/PD-L1 signaling inhibition on cognitive function and their underlying mechanisms.

This research endeavored to extensively explore the impact of blocking the PD-1/PD-L1 axis on cognitive function by integrating findings from preclinical and clinical studies. Specifically, we evaluated AD pathological and cognitive-behavioral changes in preclinical models and assessed cognitive function changes among cancer patients administered PD-1/PD-L1 inhibitors in clinical settings. In this context, this evaluation may provide valuable insights and sheds light on potential AD therapeutic targets.

## Materials and Methods

For this study, we followed the Preferred Reporting Items for Systematic Reviews and Meta-analyses (PRISMA) 2020 reporting guideline [31] (Supplementary Table 1). The study protocol was registered in the in The International Prospective Register of Systematic Reviews database (CRD42024548133). Two investigators (JY and HH) separately conducted literature search, selected study, extracted data, and assessed bias. In case of a disagreement between the two authors, two other investigators (HJC and YMY) resolved any discrepancies.

### Search strategy

We systematically searched the database of MEDLINE, Embase, CENTRAL, and Web of Science to locate pertinent preclinical and clinical studies published from their inception to April 10, 2024. We used Boolean operators with the following search keywords: “PD-1,” “PD-L1,” “immune checkpoint inhibitor,” “glia,” “astrocyte,” “amyloid-beta,” “memory,” and “cognitive function.” The specific search strategy is provided in Supplementary Table 2.

### Study Selection

If studies satisfied the following inclusion criteria, they were deemed suitable: (1) population, animals with wild type (WT) models or AD models for animal studies, and patients with any type of cancer for clinical studies; (2) intervention, either whole knock-out (KO) or hippocampal KO via virus injection for genetic deletion of PD-1 or PD-L1 in animal studies, or administration of PD-1/PD-L1 inhibitors in animal or clinical studies; (3) comparison, neither the administration of PD-1/PD-L1 inhibitors nor modulation of PD-1 signaling; (4) outcomes, pathological or cognitive-behavior changes in animal studies and alterations in cognitive function scores or the incidence of cognitive deterioration in clinical studies; and (5) study design, experimental studies for animal studies and randomized controlled trials (RCTs) with a follow-up period of up to 6 months for clinical studies. We excluded the following studies: (1) *in vitro* studies; (2) reviews, meta-analyses, non-randomized or cross-sectional studies, or case reports; (3) ongoing studies; and (4) studies that are only available as the form of abstracts or posters. The study selection was not language-restricted.

### Data Extraction

After reviewing relevant studies, the data extracted and tabulated using Microsoft Excel (2019) are as follows: first author, year of publication, study design, follow-up duration, specific information about animals and patients such as number, age, sex, mouse strain, and type of cancer, concomitant chemotherapy, information about the modulation strategy of PD-1/PD-L1, type of PD-1/PD-L1 inhibitors, including atezolizumab, avelumab, cemiplimab, dostarlimab, durvalumab, nivolumab, and pembrolizumab, prolgolimab, retifanlimab, tislelizumab, and tremelimumab, administration route and regimen of PD-1/PD-L1 inhibitors, Eastern Cooperative Oncology Group (ECOG) performance status, cancer stage, cognitive function measurement scales, and outcome details. Data from the figure outcomes were extracted by using WebPlotDigitizer 4.7 (https://apps.automeris.io/wpd/) on highly magnified images.

### Study Outcomes

Preclinical studies have evaluated both pathological and cognitive-behavioral outcomes. Pathological outcomes included changes in the area of Aβ plaques, phosphorylated tau levels, and glial fibrillary acidic protein (GFAP) intensity. The Aβ plaque deposition in the hippocampus was measured using immunohistochemistry (IHC). Phosphorylated tau levels at the Thr231 residue were quantified as the immunoreactive area by Western blotting or as a percentage of immunoreactivity by IHC. Furthermore, GFAP intensity was measured through IHC, indicating astrocytic activation. Cognitive-behavioral outcomes included changes in learning and memory functions, assessed through related tests including Radial Arm Water Maze (RAWM), Novel Object Recognition (NOR), Morris Water Maze (MWM), and Y-maze. The MWM measures the latency time (in seconds) required to reach the platform. The NOR test assesses recognition memory through the calculating the discrimination index, which indicates the proportion of time dedicated to exploring the novel object compared to the total exploration time. The RAWM evaluates spatial learning and memory based on the number of errors. Additionally, the Y-maze test evaluates spontaneous alternation, which is associated with spatial working memory, and exploratory behavior.

In clinical studies, cognitive function outcomes were assessed based on changes in cognitive function scores, and the risk of cognitive deterioration was determined through the cognitive function domain of the European Organization for Research and Treatment of Cancer QoL Questionnaire Core 30 (QLQ-C30). The domain involves the use of a 5-point Likert scale and scores range from 0 to 100. A reduction in score of ≥ 10 points from baseline, which is generally considered clinically meaningful [32], characterized cognitive deterioration,

### Statistical Analysis

Employing the meta-package in R software (version 4.3.1; R Core team 2021), all statistical analyses were conducted. For preclinical studies, the pooled standardized mean difference (SMD) with 95% confidence intervals (CIs) of pathological and cognitive-behavioral changes was calculated using the inverse variance method, considering outcomes reported across various measurement methods with different scales.

For clinical studies involving patients with cancer, the net change in scores, represented as the least square mean change in the cognitive function score from the baseline between PD-1/PD-L1 inhibitor users and non-users, were pooled. The effect size is represented as the mean difference (MD) with a 95% CIs. Moreover, the odds ratios (ORs) of cognitive deterioration in patients with cancer administered PD-1/PD-L1 inhibitors compared with untreated individuals, were pooled. Both MD and OR with 95% CIs were derived using the inverse variance method [33]. The chi-square test (*P* value <0.1) and inconsistency statistics (*I^2^)* were utilized to evaluate the heterogeneity among studies. When significant heterogeneity, as indicated by *I^2^* >50%, was observed, a random-effects model was employed to mitigate potential bias. Conversely, a fixed-effects model was utilized in case where there was no significant heterogeneity [34].

Subgroup analysis of the clinical studies was performed based on the median age, follow-up duration, and treatment regimens. To evaluate the impact of age on cognitive function, an analysis based on age was conducted using a median age of 60 years as the cutoff. Follow-up duration was categorized into three groups: up to 2 months, 3–4 months, and 5–6 months. The therapeutic protocols were categorized as either PD-1/PD-L1 inhibitor monotherapy or PD-1/PD-L1 inhibitor in combination with other chemotherapy regimens. Furthermore, to ensure the reliability of the overall pooled results, sensitivity analyses were performed, including leave-one-out method, sample size analysis, publication year analysis. Studies identified as low quality in the bias assessment were excluded.

### Quality Assessment

Employing the Systematic Review Center for Laboratory Animal Experimentation (SYRCLE) tool, a quality assessment of all involved preclinical studies was conducted [35]. The SYRCLE includes 10 domains as follows: (1) sequence generation, (2) baseline characteristics, (3) allocation concealment, (4) random housing, (5) blinding investigators, (6) random outcome assessment, (7) blinding outcome assessor, (8) incomplete outcome data, (9) selective outcome reporting, and (10) other sources of bias, with the risk of bias responses categorized as low, high, and unclear.

The Risk of Bias 2 (ROB 2) tool was implemented to conduct quality assessments of clinical studies, specifically designed for the rigorous evaluation of RCTs [36]. This tool comprises five domains: (1) randomization process, (2) deviation from intended interventions, (3) missing outcome data, (4) outcome measurement, and (5) selection of reported results. The overall risk of bias, as determined by the ROB 2, was classified as low, high, or some concern based on the five domains of bias.

## Results

### Study Selection

Presented as Figure 1, the flow diagram demonstrates selection of the eligible study in accordance with the PRISMA guidelines. After removing duplicates, 2 453 articles were reviewed by evaluating their titles and abstracts, resulting in 1 176 remaining articles. Following a full-text evaluation for eligibility, 1 143 articles were excluded. Finally, 12 animal studies [14, 19, 22, 37–45] and 21 RCTs [26–30, 46–61] were selected for meta-analysis.

**Figure 1.**
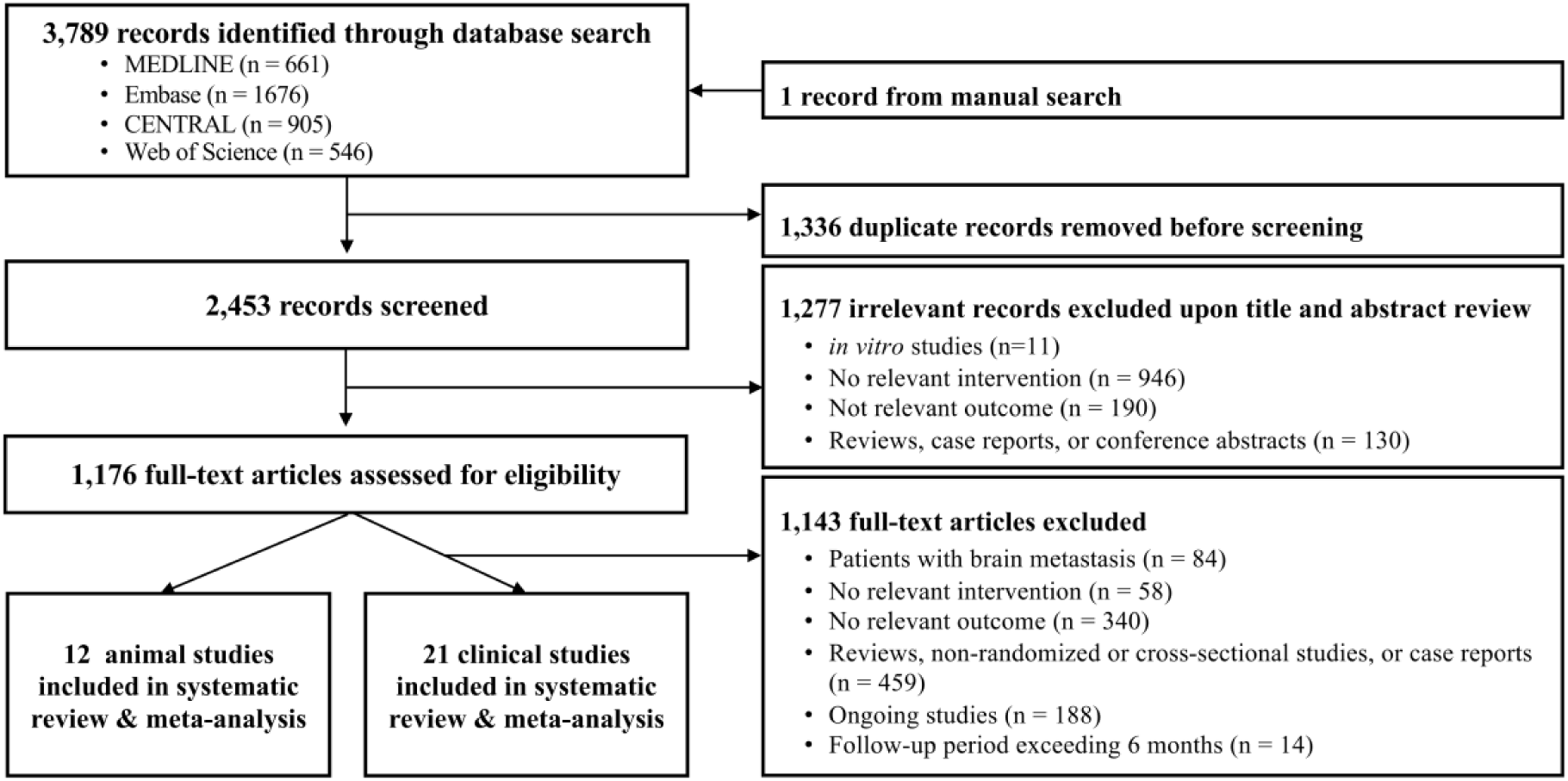
Flow chart of study selection process.

### Study Characteristics of Preclinical Studies

Among the 12 studies, PD-1 signaling was modulated in nine studies using an AD mouse model and in five studies using a disease-naive WT mouse model, namely C57BL/6. The detailed characteristics of all involved preclinical studies are presented in Supplementary Tables 3 and 4.

Among the studies using an AD mouse model, seven employed PD-1/PD-L1 antibody treatment, whereas one utilized the genetic modulation of PD-1 to inhibit PD-1 signaling. In one study, both antibody treatment and genetic deletion were used to inhibition. In both studies, PD-1 was knocked out using the genetic deletion method. Among the studies that used antibody treatment, five used anti-PD-1, and five used anti-PD-L1 antibodies. In addition, seven studies administered PD-1/PD-L1 antibody via intraperitoneal (IP) injection, whereas one study administered it via intracerebroventricular (ICV) injection. The following AD mouse models were used in the involved studies: 5XFAD in four studies, APP/PS1 in four studies, DM-Tau in two studies, JNPLT3 in one study, and Aβ insult in one study. Multiple behavioral and pathology-based tests were used in eight and six studies, respectively, to assess changes in cognitive function in AD mouse models with PD-1/PD-L1 pathway inhibition. The cognitive-behavioral tests and pathological features used in each study are described in Supplementary Table 3. Three studies involved RAWM, three involved MWM, three involved NOR, and three involved the Y-maze test. Furthermore, among the studies examining the representative pathological features of AD, five studies investigated changes in hippocampal Aβ plaque, two studies examined phosphorylated Tau levels at Thr231, and two studies assessed GFAP immunoreactivity.

Furthermore, studies using a WT mouse model showed the inhibition of PD-1 signaling through antibody treatment in one study and genetic deletion in three studies. Both antibody treatment and genetic deletion have been used in one study. Two studies employed PD-1 KO for the genetic deletion method, whereas the remaining two employed PD-L1 KO. Antibodies were administered via IP or ICV injection. Among these studies involving antibody treatment, two targeted PD-1, and one targeted PD-L1 antibodies. All included studies using the WT mouse model used behavioral tests to assess changes in cognitive function due to PD-1/PD-L1 pathway inhibition. All five studies that examined behavior using the WT mouse model conducted the MWM, while one additionally conducted the NOR.

### Study Characteristics of Clinical Studies

Supplementary Table 5 summarizes the characteristics of the included RCTs. Per study included a diverse number of participants (range, 36–512), with a total of 9 826 individuals. The median follow-up duration was 12 weeks (range, 6–24 weeks). The most frequently assessed PD-1/PD-L1 inhibitor was pembrolizumab, which was studied in 14 trials as monotherapy and in four trials as combination therapy. Monotherapy with nivolumab or cemiplimab was used in the remaining two studies. Most studies compared PD-1/PD-L1 inhibitors with other chemotherapies, while three studies assessed them against a placebo. All included studies used the cognitive function domain of the QLQ-C30 to evaluate changes in cognitive function.

Supplementary Table 6 summarizes the baseline characteristics of the study population, including a median age ranging from 35 to 67 years. Regarding the ECOG performance status score, most participants (≥99%) were distributed with a score of ≤1. Among the 21 studies, men comprised for more than half of the participants in 15 studies, while four studies predominantly included women due to specific cancer types.

### Data analysis of preclinical studies

All pooled meta-analyses showed the pathological and cognitive-behavioral effects of PD-1 signaling inhibition (Figure 2 and Supplementary Figure 1, 2).

**Figure 2.**
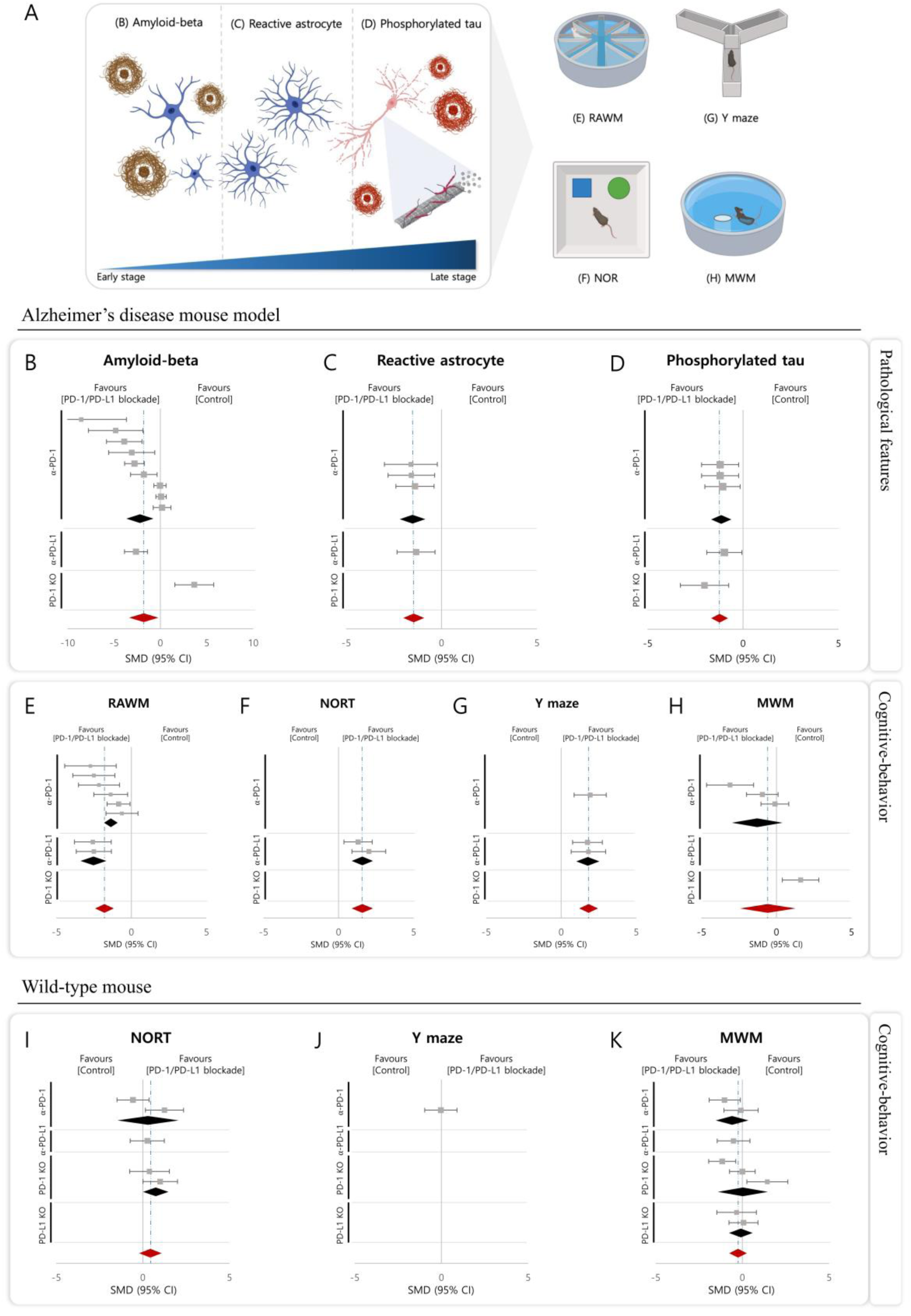
Pathological and cognitive behavioral change in mouse models blocked by PD-1/PD-L1 signaling. (A) A schematic representation of the main hallmarks of AD pathological changes and the behavioral tests used to evaluate cognitive function of mice, (B-H) Simplified forest plot of pathological and cognitive behavioral change in Alzheimer’s disease mouse model blocked by PD-1/PD-L1 signaling, (B) Amyloid-beta, (C) Reactive astrocytes, (D) Phosphorylated Tau, (E) Radical arm water maze, (F) Novel object recognition test, (G) Y maze, (H) Morris water maze of AD mouse model. (I-K) Simplified forest plot of cognitive behavioral change in wild-type mouse model blocked by PD-1/PD-L1 signaling, (I) Novel object recognition test, (J) Y maze, (K) Morris water maze of wild type mouse model. Schematic created with BioRender.com.

### Blockade of the PD-1/PD-L1 axis reduced Aβ plaques

In order to evaluate the impact of blocking PD-1/PD-L1 axis on amyloid pathology of AD mouse models, we extracted the data that measured the area covered by the Aβ plaques. PD-1/PD-L1 modulation significantly decreased the area occupied by Aβ plaques in the AD mouse model, compared to the control group (SMD = −1.80, 95% CI −3.39 to −0.22, *I^2^* = 89%, p <0.001) (Figure 2B & Supplementary Figure 1A). PD-1/PD-L1 signaling modulation by IP injection of either anti-PD-1 or anti-PD-L1 antibody was significantly associated with reduction of Aβ plaques, consistent with the overall outcome (anti-PD-1 antibody treatment: SMD = −2.17, 95% CI −3.64 to −0.71, *I^2^* = 87%; anti-PD-L1 antibody treatment: SMD = −2.63, 95% CI −3.86 to −1.39). However, in a study in which PD-1 genetic deletion was used, a different modulation method than IP administration of PD-1 antibody, an increase in Aβ plaque area was observed (SMD = 3.65, 95% CI 1.56 to 5.75). These pooled SMDs suggest that blocking PD-1/PD-L1 axis has a beneficial effect on amyloid pathology in AD.

### Blockade of the PD-1/PD-L1 axis mitigated astrocyte reactivity

Previously noted, Aβ deposits induce reactive astrocytes. Thus, we investigated the impact of blocking PD-1 signaling on reactive astrocytes by extracting and comparing the data on astrocyte reactivity (GFAP intensity measurements). Astrocyte reactivity was substantially alleviated in the group where the PD-1/PD-L1 axis was blocked compared to the control group (SMD =-1.45, 95% CI −2.01 to-0.89, *I^2^* = 0%, p <0.79) (Figure 2C & Supplementary Figure 1B). Subgroup analysis by antibody type, including anti-PD-1 and anti-PD-L1 antibodies, revealed a significant reduction in astrocyte reactivity following IP injection in an AD mouse model, which was consistent with the results of the overall analysis (anti-PD-1 antibody treatment: SMD = −1.50, 95% CI −2.18 to −0.82, *I^2^* = 0%; anti-PD-L1 antibody treatment: SMD = −1.45, 95% CI −2.01 to −0.89). This analysis showed that blocking PD-1 signaling reduced astrocyte reactivity, accompanied by the alleviation of Aβ deposits.

### Blockade of the PD-1/PD-L1 axis alleviated tau phosphorylation

Aβ-triggered reactive astrocytes cause the phosphorylation of tau protein. Therefore, we analyzed the data measuring the effect of PD-1 signaling modulation on the tau phosphorylation at the Thr231, a site showing strong relevance in AD progression [62], in AD mouse models. Blocking PD-1 signaling led to a significant reduction in the level of phosphorylated tau in the AD mouse model. (SMD= −1.22, 95% CI −1.66 to −0.78, *I^2^* = 0%) (Fig 2D & Supplementary Figure 1C). All subgroup analyses conducted based on different methods of PD-1/PD-L1 signaling modulation, through antibody intervention or genetic deletion, consistently showed a reduction in tau phosphorylation, in line with the overall analysis (anti-PD-1 antibody treatment: SMD = −1.50, 95% CI −2.18 to −0.82, *I ^2^*= 0%; anti-PD-L1 antibody treatment: SMD = −1.45, 95% CI −2.01 to −0.89; PD-1 genetic deletion: SMD = −2.02, 95% CI −3.28.18 to −0.75). The overall effect of PD-1/PD-L1 axis modulation on the tau pathology showed that PD-1 signaling inhibition mitigates AD pathological features, including amyloid pathology and reactive astrocytes.

### Blockade of the PD-1/PD-L1 axis affected cognitive function in an AD mouse model

The behavioral tests using RAWM, NOR, and Y maze test showed that blocking PD-1 signaling significantly improved the cognitive function in AD mouse models (RAWM: SMD = −1.79, 95% CI −2.41 to 1.17; *I^2^* = 55%, NORT: SMD = 1.60, 95% CI 0.88 to 2.32, *I^2^* = 0%; Y maze: SMD =1.83, 95% CI 1.21 to 2.44, *I^2^* = 0%) (Figure 2E–H & Supplementary Figure 1D–G). All studies included in the analysis involved antibody IP injection to inhibit the PD-1 signaling. Moreover, no significant variation was detected in the impact of antibody treatment on cognitive functions based on the type of antibody used (anti-PD-1 or anti-PD-L1 antibody).

However, the overall effect of blocking PD-1 signaling on the MWM behavioral test in the AD mouse model was not significant compared to the control group (SMD = −0.58, 95% CI −2.43 to 1.26, *I^2^* = 87%) (Figure 2H & Supplementary Figure 1G). The stratified analysis revealed contradictory patterns in the relationship between PD-1 signaling inhibition and latency time in the MWM, depending on the method used for signaling modulation. In one study, modulation of the PD-1 signaling through genetic deletion of PD-1 led to an increase in latency time in the MWM, indicating cognitive decline (SMD = 1.61, 95% CI 0.39 to 2.82). In contrast, the blockade through the injection of an anti-PD-1 antibody was associated with a decrease in latency time in the MWM, suggesting cognitive improvement, which aligns with the overall observed directional effect. (SMD = −1.28, 95% CI −2.94 to 0.39, *I^2^* = 81%).

### Blockade of the PD-1/PD-L1 axis showed no significant impact on cognitive function in a WT mouse model

Figure 2 I–K & Supplementary Figure 2 depicts the outcomes of a meta-analysis evaluating the effects of PD-1/PD-L1 modulation through behavioral tests in a WT mouse model. The results of behavioral tests using MWM, Y maze and NOR indicated no significant correlation between PD-1 signaling modulation and cognitive function in the WT mouse model (MWM: SMD = −0.25, 95% CI −0.76 to 0.26, *I^2^* = 60%; Y maze: SMD = −0.03, 95% CI −0.96 to 0.90; NOR: SMD = 0.44, 95% CI −0.21 to 1.08, *I^2^* = 50%). Moreover, no significant difference was observed in the association between the PD-1 signaling blockade and cognitive function assessed through behavioral tests across different methods of PD-1/PD-L1 axis modulation (antibody treatment or genetic deletion) and types of the targeted modulators (PD-1 or PD-L1).

### Data analysis of clinical studies

A meta-analysis of MDs, including 19 studies with 8 550 participants, indicated a significantly favorable impact of PD-1/PD-L1 inhibitors on the change in cognitive function scores, compared to other chemotherapies or placebo (MD = 1.88, 95% CI 0.97 to 2.78, *I^2^*= 23%). Furthermore, pooled ORs from 13 studies involving 5 410 participants showed a lower risk of cognitive deterioration among PD-1/PD-L1 inhibitor users relative to non-users (OR = 0.83, 95% CI 0.74 to 0.94, *I^2^* = 0%) (Figure 3). The results of the subgroup analyses based on median age, follow-up duration, and treatment regimen are presented in Table 1. Regardless of age, PD-1/PD-L1 inhibitor users experienced less cognitive function decline compared to the non-users (median age < 60 years: MD = 1.88, 95% CI 0.93 to 4.25, OR = 0.66, 95% CI 0.48 to 0.92; ≥ 60 years: MD = 1.58, 95% CI 0.50 to 2.66, OR = 0.86, 95% CI 0.76 to 0.98). While no significant impact was detected within 2 months, use for 3–4 months and 5– 6 months consistently favored cognitive function. Subgroup analysis based on the treatment regimen, primarily pembrolizumab, demonstrated a beneficial effect on cognitive function score changes across both pembrolizumab alone and in combination with chemotherapy groups. Specifically, in the pembrolizumab alone group, a significantly lower risk of cognitive deterioration was observed.

**Figure 3.**
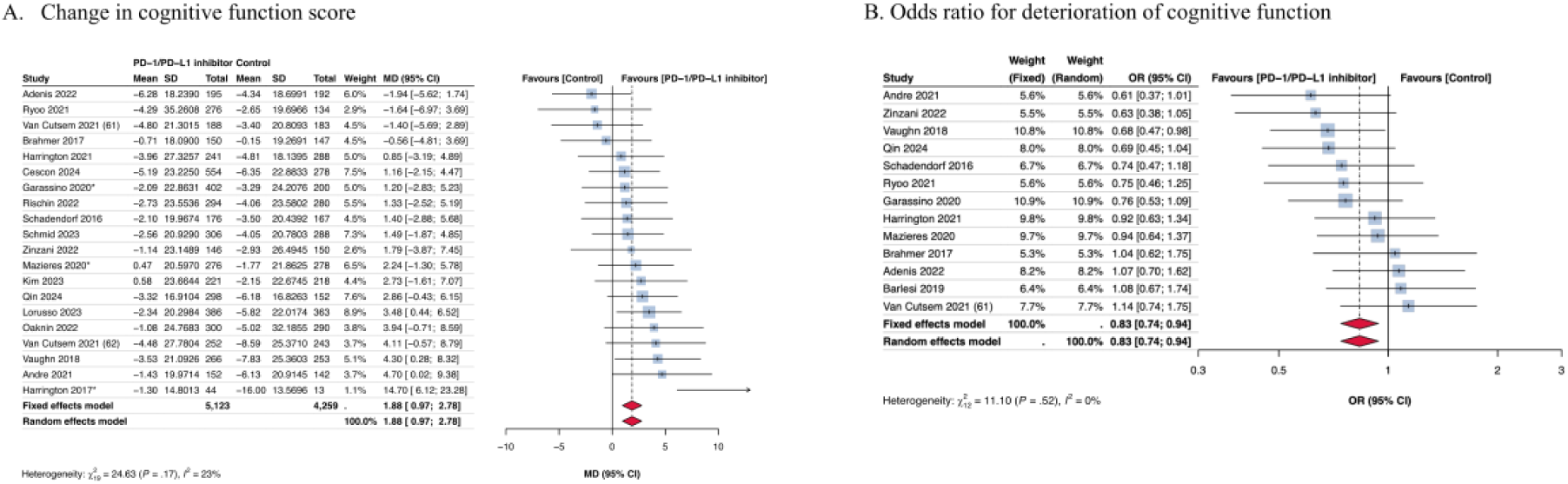
Change in cognitive function score in clinical studies. *The outcome from the longer follow-up period between two follow-up periods.

**Table 1.** Subgroup analyses of clinical studies.

| Variables | Number of studies (patients) | MD<sup>1</sup> or OR<sup>2</sup> (95% CI) | I<sup>2</sup>(%) |
| --- | --- | --- | --- |
| **Change in cognitive function score<sup>1</sup>** |  |  |  |
| *Median age* |  |  |  |
| < 60 years | 6 (2 819) | **1.88 (0.93–4.25)** | 46 |
| ≥ 60 years | 14 (6 563) | **1.58 (0.50–2.66)** | 10 |
| *Follow-up duration* |  |  |  |
| ~2 months | 5 (2 239) | 1.47 (-0.26–3.21) | 47 |
| 3-4 months | 11 (4 901) | **1.72 (0.58–2.86)** | 42 |
| 5-6 months | 6 (2 638) | **2.70 (0.92–4.47)** | 0 |
| *Treatment regimen* |  |  |  |
| Pembrolizumab alone | 13 (5 559) | **1.34 (0.20–2.48)** | 7 |
| Pembrolizumab with chemotherapy | 5 (2 874) | **1.99 (0.51–3.46)** | 0 |
| **Deterioration of cognitive function<sup>2</sup>** |  |  |  |
| *Median age* |  |  |  |
| < 60 years | 2 (373) | **0.66 (0.48–0.92)** | 0 |
| ≥ 60 years | 11 (4 664) | **0.86 (0.76–0.98)** | 0 |
| *Follow-up duration* |  |  |  |
| ~2 months | 2 (941) | 0.94 (0.71–1.24) | 0 |
| 3-4 months | 9 (3 879) | **0.85 (0.74–0.98)** | 0 |
| 5-6 months | 4 (1 746) | **0.76 (0.61–0.93)** | 0 |
| *Treatment regimen* |  |  |  |
| Pembrolizumab alone | 11 (4 254) | **0.83 (0.73–0.95)** | 5 |
| Pembrolizumab with chemotherapy | 2 (1 156) | 0.84 (0.65–1.09) | 0 |

Abbreviations: CI, confidence interval; MD, mean difference; OR, odds ratio

### Risk of Bias Assessments

Using the SYRCLE tool, assessment of bias in preclinical studies revealed that half of the studies exhibited a high risk of bias, while the other half were at an unclear risk across all domains. (Supplementary Figure 3). The domains most susceptible to bias risk were as follows: (1) selection bias, with all studies not reporting on allocation concealment; (2) performance bias, with all studies not reporting random housing but 75% of studies reporting blinded interventions; and (3) reporting bias, with 33.3% of studies showing a high risk of bias regarding selective outcome reporting. However, the detection domains reported that all studies conducted random and blinded outcome assessments, indicating clear and comprehensive reporting of outcome data.

In the clinical studies, the bias assessment using the ROB2 tool revealed that more than half of the included studies exhibited some bias, as shown in Supplementary Figure 4. This was primarily attributed to missing data owing to patient dropout, a characteristic of cancer clinical trials. Additionally, 66.7% of the studies demonstrated some concern for the risk of bias in Domain 1, while 61.9% in Domain 4, owing to the open-label and patient-reported outcome design of the studies.

## Discussion

We comprehensively evaluated the impact of PD-1/PD-L1 axis modulation on AD pathology and cognitive function via this systematic review and meta-analysis of preclinical and clinical studies. Our analysis revealed that administering anti-PD-1/PD-L1 antibodies significantly reduced the pathological hallmarks in AD mouse models, indicating its protective effect in AD pathology. Moreover, behavioral tests in AD mouse models demonstrated improvements in cognitive function, especially in the RAWM, NOR, and Y-maze tests, whereas no significance was detected in the WT mouse model. Additionally, clinical studies in cancer patients have suggested that PD-1/PD-L1 inhibitors may decrease the risk of cognitive decline compared with conventional chemotherapies, highlighting their potential benefits beyond cancer treatment.

We suggest a hypothetical mechanism for the effects of the PD-1 signaling inhibition in AD, as described below. Systemic injection of PD-1/PD-L1 inhibitors boosts the immune response, exacerbating the activation of CD8^+^ T cells and PD-1^+^ macrophages and the immunosuppression of PD-1^+^ CD4^+^ Treg cells [63, 64]. These inhibitors increase the infiltration of PD-1^+^ CD4^+^ Tregs and PD-1^+^ macrophages, but not of CD8^+^PD-1^+^T cells, through BBB disruption in animal models of AD [12, 40]. The immune cell infiltration into the brain parenchyma is believed to have protective effects on AD pathology, corroborating the results of our meta-analysis. First, infiltrating PD-1^+^ macrophages, referred to as MDMs, adopt a microglial-like morphology, migrate to amyloid plaques, and reduce plaque load [11, 65, 66]. MDM infiltration has been demonstrated to reduce pathological features of AD and alleviate cognitive decline [14, 39, 40, 43]. Second, activated Treg cells suppress the responses of activated glial cells, resulting in alleviation of neuroinflammation and AD [67, 68]. In an AD mouse model, elevated deposition of Tregs in the brain via implantation or systemic injection of anti-PD-1 antibodies significantly alleviated neurodegeneration and cognitive impairment [12, 13]. In this regard, it can be consistently explained that PD-1/PD-L1 inhibition showed no significant beneficial effects on cognitive function in studies using PD-1 genetic deletion in an AD mouse model or in a WT mouse model, when the BBB remained intact. These previous studies align with the results of our study, further highlighting the crucial role of infiltrating immune cells in PD-1/PD-L1 signaling, leading to its beneficial effects in AD.

Overall, neuroinflammation appears to be a critical factor in cognitive function, and the results from animal model-based studies suggest that systemic immune responses influence neuroinflammation. Currently, PD-1/PD-L1 inhibitors are used exclusively for treating cancer in clinical settings; therefore, their effects on cognitive function can only be assessed in patients with cancer. However, cancer itself is one of the causes of systemic inflammation, and this inflammatory mechanism can also affect the brain, thereby influencing cognitive function [69–71]. Therefore, despite the expected cognitive decline in cancer patients, our meta-analysis of clinical studies showed that PD-1/PD-L1 inhibitor users experienced a smaller decrease in cognitive function scores compared to non-users. In addition, we observed that PD-1/PD-L1 inhibitor users had a relatively diminished incidence of cognitive deterioration, compared with non-users. Although the clinical trial participants were not patients with AD, our results aligned with the findings of preclinical studies, suggesting that neuroinflammation was influenced by the PD-1/PD-L1 pathway.

Just as brain structure and function changes with age, cognitive changes are also linked to age [72]. Nevertheless, our age-based analyses showed a consistent and significant advantageous effect on cognitive function in PD-1/PD-L1 inhibitor users, both in those with a median age of 60 years or older and in those who were younger. While we conducted an age-based analysis using a median age-based cutoff of 60 years, the diverse age ranges across studies might have influenced our results. This finding suggests the need for further studies to comprehensively elucidate the effect of PD-1/PD-L1 inhibitor use on cognitive function across limited age groups. In addition to the age, subgroup analyses considering follow-up duration and treatment regimen provided additional insights into the effects of PD-1/PD-L1 inhibitors on cognitive decline. Although the effect of PD-1/PD-L1 inhibitors did not achieve statistical significance with a 2-month duration use, a trend favoring cognitive function was observed. Considering that the time-to-response to PD-1/PD-L1 inhibitors is approximately 2 months on average, these results indicate that the cognitive outcomes may hinge on the length of time it takes for these inhibitors to exert their effects in cancer patients [73, 74]. In contrast, our analysis found a significant reduction in the risk of cognitive function decline among patients who administered PD-1/PD-L1 inhibitors for 3–4 months and 5–6 months, highlighting the potential for sustained cognitive benefits with longer-term use. However, we did not evaluate a follow-up period of over 6 months due to the increase in the mortality rate over time in cancer patients [75]. Thus, additional research is required to ascertain the effects of prolonged PD-1/PD-L1 inhibitor use on cognitive function. Furthermore, our subgroup analysis based on treatment regimens revealed that both PD-1/PD-L1 inhibitor monotherapy and combination therapy tended to reduce cognitive function decline. Given that the OR of PD-1/PD-L1 inhibitor combination therapy alone was not statistical significance, the differences between monotherapy and combination therapy in terms of the risk of cognitive deterioration may be challenging to explain. This result may be attributed to the relatively limited number of patients or the potential negative effects related to neurotoxicity and health status due to the anticancer therapy [76].

Although our study contributed significantly to understanding the impact of PD-1/PD-L1 inhibitors on cognitive function, several limitations should be acknowledged. First, the included preclinical and clinical studies were highly heterogeneous. In preclinical studies, various PD-1/PD-L1 signaling modulation methods, the types of AD mouse models, and behavioral outcomes have been employed to test cognitive decline. In addition, clinical studies that include different types of cancers can lead to variability in the response to treatment and in cognitive outcomes. Therefore, we endeavored to provide consolidated results, through sensitivity analyses, that aligned with our main findings. These analytical approaches helped account for the heterogeneity included in the study and enhanced the reliability of our conclusions. Moreover, by performing subgroups analyses, based on the modulation method in preclinical studies or follow-up duration in clinical studies, we can suggest a mechanism-based understanding of the observed results and conduct a more in-depth review. Second, each study may have potential biases. In preclinical studies, while multiple outcomes were extracted and individually weighted in the analysis from a single research study, the results all exhibited the same trend and were significantly influenced by paper bias. In addition, some subgroups defined by the modulation methods included only a single study, revealing a high risk of bias in the quality assessment. In clinical studies, the relatively high dropout rates, which are often inherent in studies involving patients with cancer, need to be considered. The high dropout rates observed across studies could result in missing data and introduce bias, potentially affecting the reliability of our results [75]. Moreover, the open-label characteristics of the included studies raise concerns about a potential reporting bias, as both patients and researchers were aware of the treatment assignment. Finally, the lack of international guidelines for the statistical analysis of health-related quality of life data may have led to inconsistencies in the assessment and interpretation of cognitive function-based outcomes across studies [77]. These limitations underscore the need for a careful interpretation of our study, suggest the requirement of further research to address these methodological challenges, confirm the robustness of our results.

In conclusion, our study unveiled a novel therapeutic avenue for AD and proposed the potential drug repositioning of PD-1/PD-L1 inhibitors, originally used as anti-cancer drugs. Moreover, it underscored the importance of advancing neuroscientific exploration into how PD-1/PD-L1 signaling impacts AD and cognitive decline.

## Acknowledgements

This work was supported by the Yonsei University Research Fund of 2024 (2024-22-0131 to H.C.) and by Basic Science Research Program through the National Research Foundation of Korea(NRF) funded by the Ministry of Education(2018R1A6A1A03023718).

## Additional information

Supplementary information is available at Molecular Psychiatry’s website.

## References

1 Duyckaerts C, Delatour B, Potier MC. Classification and basic pathology of Alzheimer disease. Acta Neuropathol. 2009;118(1):5–36.

2 Masters CL, Bateman R, Blennow K, Rowe CC, Sperling RA, Cummings JL. Alzheimer’s disease. Nat Rev Dis Primers. 2015;1:15056.

3 Allaman I, Gavillet M, Bélanger M, Laroche T, Viertl D, Lashuel HA, et al. Amyloid-beta aggregates cause alterations of astrocytic metabolic phenotype: impact on neuronal viability. J Neurosci. 2010;30(9):3326–38.

4 Chun H, Lee CJ. Reactive astrocytes in Alzheimer’s disease: A double-edged sword. Neuroscience Research. 2018;126:44–52.

5 Orre M, Kamphuis W, Osborn LM, Jansen AHP, Kooijman L, Bossers K, et al. Isolation of glia from Alzheimer’s mice reveals inflammation and dysfunction. Neurobiol Aging. 2014;35(12):2746–60.

6 Bellaver B, Povala G, Ferreira PCL, Ferrari-Souza JP, Leffa DT, Lussier FZ, et al. Astrocyte reactivity influences amyloid-β effects on tau pathology in preclinical Alzheimer’s disease. Nature Medicine. 2023;29(7):1775–81.

7 Chun H, Im H, Kang YJ, Kim Y, Shin JH, Won W, et al. Severe reactive astrocytes precipitate pathological hallmarks of Alzheimer’s disease via H(2)O(2)(-) production. Nat Neurosci. 2020;23(12):1555–66.

8 Cullen NC, Mälarstig An, Stomrud E, Hansson O, Mattsson-Carlgren N. Accelerated inflammatory aging in Alzheimer’s disease and its relation to amyloid, tau, and cognition. Scientific Reports. 2021;11(1):1965.

9 Ballatore C, Lee VMY, Trojanowski JQ. Tau-mediated neurodegeneration in Alzheimer’s disease and related disorders. Nature Reviews Neuroscience. 2007;8(9):663–72.

10 Jorfi M, Park J, Hall CK, Lin C-CJ, Chen M, von Maydell D, et al. Infiltrating CD8+ T cells exacerbate Alzheimer’s disease pathology in a 3D human neuroimmune axis model. Nature Neuroscience. 2023;26(9):1489–504.

11 Muñoz-Castro C, Mejias-Ortega M, Sanchez-Mejias E, Navarro V, Trujillo-Estrada L, Jimenez S, et al. Monocyte-derived cells invade brain parenchyma and amyloid plaques in human Alzheimer’s disease hippocampus. Acta Neuropathologica Communications. 2023;11(1):31.

12 Chen X, Firulyova M, Manis M, Herz J, Smirnov I, Aladyeva E, et al. Microglia-mediated T cell infiltration drives neurodegeneration in tauopathy. Nature. 2023;615(7953):668–77.

13 Baek H, Ye M, Kang GH, Lee C, Lee G, Choi DB, et al. Neuroprotective effects of CD4+CD25+Foxp3+ regulatory T cells in a 3xTg-AD Alzheimer’s disease model. Oncotarget. 2016;7(43):69347–57.

14 Dvir-Szternfeld R, Castellani G, Arad M, Cahalon L, Colaiuta SP, Keren-Shaul H, et al. Alzheimer’s disease modification mediated by bone marrow-derived macrophages via a TREM2-independent pathway in mouse model of amyloidosis. Nature Aging. 2022;2(1):60–73.

15 Agata Y, Kawasaki A, Nishimura H, Ishida Y, Tsubata T, Yagita H, et al. Expression of the PD-1 antigen on the surface of stimulated mouse T and B lymphocytes. Int Immunol. 1996;8(5):765–72.

16 Bardhan K, Anagnostou T, Boussiotis VA. The PD1:PD-L1/2 Pathway from Discovery to Clinical Implementation. Front Immunol. 2016;7:550.

17 Zhao J, Ji R-R. Anti-PD-1 treatment as a neurotherapy to enhance neuronal excitability, synaptic plasticity and memory. bioRxiv. 2019:870600.

18 Wang Z, Jiang C, He Q, Matsuda M, Han Q, Wang K, et al. Anti-PD-1 treatment impairs opioid antinociception in rodents and nonhuman primates. Sci Transl Med. 2020;12(531).

19 Zhao J, Bang S, Furutani K, McGinnis A, Jiang C, Roberts A, et al. PD-L1/PD-1 checkpoint pathway regulates hippocampal neuronal excitability and learning and memory behavior. Neuron. 2023;111(17):2709–26.e9.

20 Zhao J, Roberts A, Wang Z, Savage J, Ji R-R. Emerging Role of PD-1 in the Central Nervous System and Brain Diseases. Neuroscience Bulletin. 2021;37(8):1188–202.

21 Schachtele SJ, Hu S, Sheng WS, Mutnal MB, Lokensgard JR. Glial cells suppress postencephalitic CD8+ T lymphocytes through PD-L1. Glia. 2014;62(10):1582–94.

22 Kummer MP, Ising C, Kummer C, Sarlus H, Griep A, Vieira-Saecker A, et al. Microglial PD-1 stimulation by astrocytic PD-L1 suppresses neuroinflammation and Alzheimer’s disease pathology. The EMBO Journal. 2021;40(24):e108662.

23 Linnerbauer M, Beyer T, Nirschl L, Farrenkopf D, Lößlein L, Vandrey O, et al. PD-L1 positive astrocytes attenuate inflammatory functions of PD-1 positive microglia in models of autoimmune neuroinflammation. Nature Communications. 2023;14(1):5555.

24 Sun L, Zhang L, Yu J, Zhang Y, Pang X, Ma C, et al. Clinical efficacy and safety of anti-PD-1/PD-L1 inhibitors for the treatment of advanced or metastatic cancer: a systematic review and meta-analysis. Sci Rep. 2020;10(1):2083.

25 Wankhede D, Hofman P, Grover S. PD-1/PD-L1 inhibitors in treatment-naïve, advanced non-small cell lung cancer patients with & 1% PD-L1 expression: a meta-analysis of randomized controlled trials. J Cancer Res Clin Oncol. 2023;149(5):2179–89.

26 Vaughn DJ, Bellmunt J, Fradet Y, Lee JL, Fong L, Vogelzang NJ, et al. Health-Related Quality-of-Life Analysis From KEYNOTE-045: A Phase III Study of Pembrolizumab Versus Chemotherapy for Previously Treated Advanced Urothelial Cancer. J Clin Oncol. 2018;36(16):1579–87.

27 Andre T, Amonkar M, Norquist JM, Shiu KK, Kim TW, Jensen BV, et al. Health-related quality of life in patients with microsatellite instability-high or mismatch repair deficient metastatic colorectal cancer treated with first-line pembrolizumab versus chemotherapy (KEYNOTE-177): an open-label, randomised, phase 3 trial. Lancet Oncol. 2021;22(5):665–77.

28 Adenis A, Kulkarni AS, Girotto GC, de la Fouchardiere C, Senellart H, van Laarhoven HWM, et al. Impact of Pembrolizumab Versus Chemotherapy as Second-Line Therapy for Advanced Esophageal Cancer on Health-Related Quality of Life in KEYNOTE-181. Journal of clinical oncology. 2022;40(4):382-91.

29 Ryoo BY, Merle P, Kulkarni AS, Cheng AL, Bouattour M, Lim HY, et al. Health-related quality-of-life impact of pembrolizumab versus best supportive care in previously systemically treated patients with advanced hepatocellular carcinoma: KEYNOTE-240. Cancer. 2021;127(6):865-74.

30 Brahmer JR, Rodriguez-Abreu D, Robinson AG, Hui R, Csoszi T, Fulop A, et al. Health-related quality-of-life results for pembrolizumab versus chemotherapy in advanced, PD-L1-positive NSCLC (KEYNOTE-024): a multicentre, international, randomised, open-label phase 3 trial. The lancet Oncology. 2017;18(12):1600–09.

31 Page MJ, McKenzie JE, Bossuyt PM, Boutron I, Hoffmann TC, Mulrow CD, et al. The PRISMA 2020 statement: an updated guideline for reporting systematic reviews. Bmj. 2021;372:n71.

32 Osoba D, Rodrigues G, Myles J, Zee B, Pater J. Interpreting the significance of changes in health-related quality-of-life scores. J Clin Oncol. 1998;16(1):139–44.

33 Hahn J, Jeon J, Geum MJ, Lee HW, Shin J, Chung WY, et al. Intracoronary versus intravenous glycoprotein IIb/IIIa inhibitors during primary percutaneous coronary intervention in patients with STEMI: a systematic review and meta-analysis. Thromb J. 2023;21(1):76.

34 Higgins JP, Thompson SG. Quantifying heterogeneity in a meta-analysis. Stat Med. 2002;21(11):1539–58.

35 Hooijmans CR, Rovers MM, de Vries RB, Leenaars M, Ritskes-Hoitinga M, Langendam MW. SYRCLE’s risk of bias tool for animal studies. BMC Med Res Methodol. 2014;14:43.

36 Sterne JAC, Savović J, Page MJ, Elbers RG, Blencowe NS, Boutron I, et al. RoB 2: a revised tool for assessing risk of bias in randomised trials. Bmj. 2019;366:l4898.

37 Zou Y, Gan CL, Xin Z, Zhang HT, Zhang Q, Lee TH, et al. Programmed Cell Death Protein 1 Blockade Reduces Glycogen Synthase Kinase 3β Activity and Tau Hyperphosphorylation in Alzheimer’s Disease Mouse Models. Front Cell Dev Biol. 2021;9:769229.

38 Xing Z, Zuo Z, Hu D, Zheng X, Wang X, Yuan L, et al. Influenza vaccine combined with moderate-dose PD1 blockade reduces amyloid-β accumulation and improves cognition in APP/PS1 mice. Brain Behav Immun. 2021;91:128–41.

39 Ben-Yehuda H, Arad M, Peralta Ramos JM, Sharon E, Castellani G, Ferrera S, et al. Key role of the CCR2-CCL2 axis in disease modification in a mouse model of tauopathy. Molecular neurodegeneration. 2021;16(1):1–19.

40 Rosenzweig N, Dvir-Szternfeld R, Tsitsou-Kampeli A, Keren-Shaul H, Ben-Yehuda H, Weill-Raynal P, et al. PD-1/PD-L1 checkpoint blockade harnesses monocyte-derived macrophages to combat cognitive impairment in a tauopathy mouse model. Nature Communications. 2019;10(1):465.

41 Lin Y, Rajamohamedsait HB, Sandusky-Beltran LA, Gamallo-Lana B, Mar A, Sigurdsson EM. Chronic PD-1 Checkpoint Blockade Does Not Affect Cognition or Promote Tau Clearance in a Tauopathy Mouse Model. Front Aging Neurosci. 2019;11:377.

42 Karl F, Colaço MBN, Schulte A, Sommer C, Üçeyler N. Affective and cognitive behavior is not altered by chronic constriction injury in B7-H1 deficient and wildtype mice. BMC Neurosci. 2019;20(1):16.

43 Baruch K, Deczkowska A, Rosenzweig N, Tsitsou-Kampeli A, Sharif AM, Matcovitch-Natan O, et al. PD-1 immune checkpoint blockade reduces pathology and improves memory in mouse models of Alzheimer’s disease. Nature Medicine. 2016;22(2):135–37.

44 Latta-Mahieu M, Elmer B, Bretteville A, Wang Y, Lopez-Grancha M, Goniot P, et al. Systemic immune-checkpoint blockade with anti-PD1 antibodies does not alter cerebral amyloid-β burden in several amyloid transgenic mouse models. Glia. 2018;66(3):492–504.

45 Karl F, Grießhammer A, Üçeyler N, Sommer C. Differential Impact of miR-21 on Pain and Associated Affective and Cognitive Behavior after Spared Nerve Injury in B7-H1 ko Mouse. Front Mol Neurosci. 2017;10:219.

46 Barlesi F, Garon EB, Kim DW, Felip E, Han JY, Kim JH, et al. Health-Related Quality of Life in KEYNOTE-010: a Phase II/III Study of Pembrolizumab Versus Docetaxel in Patients With Previously Treated Advanced, Programmed Death Ligand 1-Expressing NSCLC. Journal of thoracic oncology. 2019;14(5):793–801.

47 Garassino MC, Gadgeel S, Esteban E, Felip E, Speranza G, Domine M, et al. Patient-reported outcomes following pembrolizumab or placebo plus pemetrexed and platinum in patients with previously untreated, metastatic, non-squamous non-small-cell lung cancer (KEYNOTE-189): a multicentre, double-blind, randomised, placebo-controlled, phase 3 trial. The lancet Oncology. 2020;21(3):387–97.

48 Harrington KJ, Ferris RL, Blumenschein G, Colevas AD, Fayette J, Licitra L, et al. Nivolumab versus standard, single-agent therapy of investigator’s choice in recurrent or metastatic squamous cell carcinoma of the head and neck (CheckMate 141): health-related quality-of-life results from a randomised, phase 3 trial. The lancet Oncology. 2017;18(8):1104–15.

49 Harrington KJ, Soulières D, Le Tourneau C, Dinis J, Licitra LF, Ahn MJ, et al. Quality of Life With Pembrolizumab for Recurrent and/or Metastatic Head and Neck Squamous Cell Carcinoma: KEYNOTE-040. J Natl Cancer Inst. 2021;113(2):171–81.

50 Kim HR, Awad MM, Navarro A, Gottfried M, Peters S, Csőszi T, et al. Patient-Reported Health-Related Quality of Life in KEYNOTE-604: Pembrolizumab or Placebo Added to Etoposide and Platinum as First-Line Therapy for Extensive-Stage SCLC. JTO Clin Res Rep. 2023;4(11):100572.

51 Lorusso D, Colombo N, Herraez AC, Santin AD, Colomba E, Miller DS, et al. Health-Related Quality of Life in Patients With Advanced Endometrial Cancer Treated With Lenvatinib Plus Pembrolizumab or Treatment of Physician’s Choice. European journal of cancer (Oxford, England : 1990). 2023;186:172–84.

52 Mazieres J, Kowalski D, Luft A, Vicente D, Tafreshi A, Gümüş M, et al. Health-Related Quality of Life With Carboplatin-Paclitaxel or nab-Paclitaxel With or Without Pembrolizumab in Patients With Metastatic Squamous Non-Small-Cell Lung Cancer. Journal of clinical oncology. 2020;38(3):271–80.

53 Oaknin A, Monk BJ, Vergote I, Cristina de Melo A, Kim YM, Lisyanskaya AS, et al. EMPOWER CERVICAL-1: Effects of cemiplimab versus chemotherapy on patient-reported quality of life, functioning and symptoms among women with recurrent cervical cancer. Eur J Cancer. 2022;174:299–309.

54 Qin S, Fang W, Ren Z, Ou S, Lim HY, Zhang F, et al. A Phase 3 Study of Pembrolizumab versus Placebo for Previously Treated Patients from Asia with Hepatocellular Carcinoma: Health-Related Quality of Life Analysis from KEYNOTE-394. Liver Cancer. 2024:1–12.

55 Rischin D, Harrington KJ, Greil R, Soulières D, Tahara M, de Castro G, et al. Pembrolizumab alone or with chemotherapy for recurrent or metastatic head and neck squamous cell carcinoma: health-related quality-of-life results from KEYNOTE-048. Oral oncology. 2022;128:105815.

56 Schadendorf D, Dummer R, Hauschild A, Robert C, Hamid O, Daud A, et al. Health-related quality of life in the randomised KEYNOTE-002 study of pembrolizumab versus chemotherapy in patients with ipilimumab-refractory melanoma. European journal of cancer (Oxford, England : 1990). 2016;67:46–54.

57 Schmid P, Lipatov O, Im SA, Goncalves A, Muñoz-Couselo E, Lee KS, et al. Impact of pembrolizumab versus chemotherapy on health-related quality of life in patients with metastatic triple-negative breast cancer: results from the phase 3 randomised KEYNOTE-119 study. European journal of cancer (Oxford, England : 1990). 2023;195:113393.

58 Van Cutsem E, Amonkar M, Fuchs CS, Alsina M, Özgüroğlu M, Bang YJ, et al. Health-related quality of life in advanced gastric/gastroesophageal junction cancer with second-line pembrolizumab in KEYNOTE-061. Gastric cancer. 2021;24(6):1330–40.

59 Van Cutsem E, Valderrama A, Bang YJ, Fuchs CS, Shitara K, Janjigian YY, et al. Quality of life with first-line pembrolizumab for PD-L1epositive advanced gastric/gastroesophageal junction adenocarcinoma: results from the randomised phase III KEYNOTE-062 study. ESMO Open. 2021;6(4):11.

60 Zinzani PL, Ramchandren R, Santoro A, Paszkiewicz-Kozik E, Gasiorowski R, Johnson NA, et al. Quality-of-life analysis of pembrolizumab vs brentuximab vedotin for relapsed/refractory classical Hodgkin lymphoma. Blood advances. 2022;6(2):590–99.

61 Cescon DW, Schmid P, Rugo HS, Im SA, Md Yusof M, Gallardo C, et al. Health-related quality of life with pembrolizumab plus chemotherapy vs placebo plus chemotherapy for advanced triple-negative breast cancer: KEYNOTE-355. J Natl Cancer Inst. 2024;116(5):717–27.

62 Neddens J, Temmel M, Flunkert S, Kerschbaumer B, Hoeller C, Loeffler T, et al. Phosphorylation of different tau sites during progression of Alzheimer’s disease. Acta Neuropathologica Communications. 2018;6(1):52.

63 Kumagai S, Togashi Y, Kamada T, Sugiyama E, Nishinakamura H, Takeuchi Y, et al. The PD-1 expression balance between effector and regulatory T cells predicts the clinical efficacy of PD-1 blockade therapies. Nat Immunol. 2020;21(11):1346–58.

64 Gordon SR, Maute RL, Dulken BW, Hutter G, George BM, McCracken MN, et al. PD-1 expression by tumour-associated macrophages inhibits phagocytosis and tumour immunity. Nature. 2017;545(7655):495–99.

65 Simard AR, Soulet D, Gowing G, Julien J-P, Rivest S. Bone marrow-derived microglia play a critical role in restricting senile plaque formation in Alzheimer’s disease. Neuron. 2006;49(4):489–502.

66 Yan P, Kim K-W, Xiao Q, Ma X, Czerniewski LR, Liu H, et al. Peripheral monocyte–derived cells counter amyloid plaque pathogenesis in a mouse model of Alzheimer’s disease. The Journal of Clinical Investigation. 2022;132(11).

67 Faridar A, Vasquez M, Thome AD, Yin Z, Xuan H, Wang JH, et al. Ex vivo expanded human regulatory T cells modify neuroinflammation in a preclinical model of Alzheimer’s disease. Acta Neuropathologica Communications. 2022;10(1):144.

68 Ito M, Komai K, Mise-Omata S, Iizuka-Koga M, Noguchi Y, Kondo T, et al. Brain regulatory T cells suppress astrogliosis and potentiate neurological recovery. Nature. 2019;565(7738):246–50.

69 Oppegaard K, Harris CS, Shin J, Paul SM, Cooper BA, Chan A, et al. Cancer-related cognitive impairment is associated with perturbations in inflammatory pathways. Cytokine. 2021;148:155653.

70 Olson B, Marks DL. Pretreatment Cancer-Related Cognitive Impairment-Mechanisms and Outlook. Cancers (Basel). 2019;11(5).

71 Janelsins MC, Kesler SR, Ahles TA, Morrow GR. Prevalence, mechanisms, and management of cancer-related cognitive impairment. Int Rev Psychiatry. 2014;26(1):102–13.

72 Murman DL. The Impact of Age on Cognition. Semin Hear. 2015;36(3):111–21.

73 Inamoto T, Sato R, Matsushita Y, Uchimoto T, Nakamura KO, Komura K, et al. Optimal Time Point for Evaluation of Response to Pembrolizumab Treatment in Japanese Patients With Metastatic Urothelial Carcinoma. Cancer Diagn Progn. 2023;3(3):370–76.

74 Bironzo P, Passiglia F, Novello S. Five-year overall survival of pembrolizumab in advanced non-small cell lung cancer: another step from care to cure? Ann Transl Med. 2019;7(Suppl 6):S212.

75 Gebert P, Schindel D, Frick J, Schenk L, Grittner U. Characteristics and patient-reported outcomes associated with dropout in severely affected oncological patients: an exploratory study. BMC Med Res Methodol. 2021;21(1):77.

76 Stone JB, DeAngelis LM. Cancer-treatment-induced neurotoxicity--focus on newer treatments. Nat Rev Clin Oncol. 2016;13(2):92–105.

77 Boutros A, Bruzzone M, Tanda ET, Croce E, Arecco L, Cecchi F, et al. Health-related quality of life in cancer patients treated with immune checkpoint inhibitors in randomised controlled trials: A systematic review and meta-analysis. Eur J Cancer. 2021;159:154–66.

