## Supplementary figure for "Potential beneficial effects of PD-1/PD-L1 blockade in Alzheimer’s disease: A systematic review and meta-analysis of preclinical and clinical studies"

**Contents**

**Supplementary Table 1.** Checklist for Preferred Reporting Items for Systematic Reviews and Meta-Analyses

**Supplementary Table 2.** Search strategy

**Supplementary Table 3.** Characteristics of included preclinical studies with mouse models of Alzheimer’s disease

**Supplementary Table 4.** Characteristics of included preclinical studies with wild-type mouse models

**Supplementary Table 5.** Characteristic of included clinical studies

**Supplementary Table 6.** Baseline characteristics of the study population in clinical studies

**Supplementary Figure 1.** Detailed forest plot of pathological and cognitive behavioral changes in Alzheimer’s disease mouse model blocked by PD-1/PD-L1 signaling

**Supplementary Figure 2.** Detailed forest plot of cognitive behavioral changes in wild-type mouse model blocked by PD-1/PD-L1 signaling

**Supplementary figure 3.** Quality assessment of the risk of bias according to the Systematic Review Centre for Laboratory Animal Experimentation criteria

**Supplementary figure 4.** Quality assessments of the risk of bias according to the Risk of Bias 2 criteria

**Supplementary Figure 5.** Sensitivity analysis

Supplementary Table 1. Checklist for Preferred Reporting Items for Systematic Reviews and Meta-Analyses

| **Section/topic** | **#** | **Checklist item** | **Reported on page #** |
| --- | --- | --- | --- |
| **TITLE** |  |  |  |
| Title | 1 | Identify the report as a systematic review, meta-analysis, or both. |  |
| **ABSTRACT** |  | |  |
| Structured summary | 2 | Provide a structured summary including, as applicable: background; objectives; data sources; study eligibility criteria, participants, and interventions; study appraisal and synthesis methods; results; limitations; conclusions and implications of key findings; systematic review registration number. |  |
| **INTRODUCTION** |  |  |  |
| Rationale | 3 | Describe the rationale for the review in the context of what is already known. |  |
| Objectives | 4 | Provide an explicit statement of questions being addressed with reference to participants, interventions, comparisons, outcomes, and study design (PICOS). |  |
| **METHODS** | | | |
| Protocol and registration | 5 | Indicate if a review protocol exists, if and where it can be accessed (e.g., Web address), and, if available, provide registration information including registration number. |  |
| Eligibility criteria | 6 | Specify study characteristics (e.g., PICOS, length of follow-up) and report characteristics (e.g., years considered, language, publication status) used as criteria for eligibility, giving rationale. |  |
| Information sources | 7 | Describe all information sources (e.g., databases with dates of coverage, contact with study authors to identify additional studies) in the search and date last searched. |  |
| Search | 8 | Present full electronic search strategy for at least one database, including any limits used, such that it could be repeated. |  |
| Study selection | 9 | State the process for selecting studies (i.e., screening, eligibility, included in systematic review, and, if applicable, included in the meta-analysis). |  |
| Data collection process | 10 | Describe method of data extraction from reports (e.g., piloted forms, independently, in duplicate) and any processes for obtaining and confirming data from investigators. |  |
| Data items | 11 | List and define all variables for which data were sought (e.g., PICOS, funding sources) and any assumptions and simplifications made. |  |
| Risk of bias in individual studies | 12 | Describe methods used for assessing risk of bias of individual studies (including specification of whether this was done at the study or outcome level), and how this information is to be used in any data synthesis. |  |
| Summary measures | 13 | State the principal summary measures (e.g., risk ratio, difference in means). |  |
| Synthesis of results | 14 | Describe the methods of handling data and combining results of studies, if done, including measures of consistency (e.g., I^2^) for each meta-analysis. |  |
| Risk of bias across studies | 15 | Specify any assessment of risk of bias that may affect the cumulative evidence (e.g., publication bias, selective reporting within studies). |  |
| Additional analyses | 16 | Describe methods of additional analyses (e.g., sensitivity or subgroup analyses, meta-regression), if done, indicating which were pre-specified. |  |
| **RESULTS** | | | |
| Study selection | 17 | Give numbers of studies screened, assessed for eligibility, and included in the review, with reasons for exclusions at each stage, ideally with a flow diagram. |  |
| Study characteristics | 18 | For each study, present characteristics for which data were extracted (e.g., study size, PICOS, follow-up period) and provide the citations. |  |
| Risk of bias within studies | 19 | Present data on risk of bias of each study and, if available, any outcome level assessment (see item 12). |  |
| Results of individual studies | 20 | For all outcomes considered (benefits or harms), present, for each study: (a) simple summary data for each intervention group (b) effect estimates and confidence intervals, ideally with a forest plot. |  |
| Synthesis of results | 21 | Present results of each meta-analysis done, including confidence intervals and measures of consistency. |  |
| Risk of bias across studies | 22 | Present results of any assessment of risk of bias across studies (see Item 15). |  |
| Additional analysis | 23 | Give results of additional analyses, if done (e.g., sensitivity or subgroup analyses, meta-regression [see Item 16]). |  |
| **DISCUSSION** |  |  |  |
| Summary of evidence | 24 | Summarize the main findings including the strength of evidence for each main outcome; consider their relevance to key groups (e.g., healthcare providers, users, and policy makers). |  |
| Limitations | 25 | Discuss limitations at study and outcome level (e.g., risk of bias), and at review-level (e.g., incomplete retrieval of identified research, reporting bias). |  |
| Conclusions | 26 | Provide a general interpretation of the results in the context of other evidence, and implications for future research. |  |
| **FUNDING** |  |  |  |
| Funding | 27 | Describe sources of funding for the systematic review and other support (e.g., supply of data); role of funders for the systematic review. |  |

Supplementary Table 2. Search strategy

| **MEDLINE** | |
| --- | --- |
| #1 | "PD-1"[Title/Abstract] OR "PD1"[Title/Abstract] OR "CD279"[Title/Abstract] OR "PD-L1"[Title/Abstract] OR "PDL1"[Title/Abstract] OR "B7-H1"[Title/Abstract] OR "B7H1"[Title/Abstract] OR "CD274"[Title/Abstract] OR "CTLA-4"[Title/Abstract] OR "CTLA4"[Title/Abstract] OR "CD152"[Title/Abstract] OR "immune checkpoint"[Title/Abstract] OR "immune checkpoints"[Title/Abstract] OR "stimulatory checkpoint"[Title/Abstract] OR "stimulatory checkpoints"[Title/Abstract] OR "pd l1 inhibitor"[Title/Abstract] OR "pd l1 inhibitors"[Title/Abstract] OR "pd l1 blocker"[Title/Abstract] OR "pd l1 blockers"[Title/Abstract] OR "pd l1 suppressor"[Title/Abstract] OR ("PD-L1"[All Fields] AND "suppressors"[Title/Abstract]) OR "pdl1 inhibitor"[Title/Abstract] OR "pdl1 inhibitors"[Title/Abstract] OR ("PDL1"[All Fields] AND "blocker"[Title/Abstract]) OR ("PDL1"[All Fields] AND "blockers"[Title/Abstract]) OR ("PDL1"[All Fields] AND "suppressor"[Title/Abstract]) OR ("PDL1"[All Fields] AND "suppressors"[Title/Abstract]) OR "anti-PDL1"[Title/Abstract] OR "pd 1 inhibitor"[Title/Abstract] OR "pd 1 inhibitors"[Title/Abstract] OR "pd 1 blocker"[Title/Abstract] OR "pd 1 blockers"[Title/Abstract] OR ("PD-1"[All Fields] AND "suppressor"[Title/Abstract]) OR ("PD-1"[All Fields] AND "suppressors"[Title/Abstract]) OR "pd1 inhibitor"[Title/Abstract] OR "pd1 inhibitors"[Title/Abstract] OR "pd1 blocker"[Title/Abstract] OR ("PD1"[All Fields] AND "blockers"[Title/Abstract]) OR ("PD1"[All Fields] AND "suppressor"[Title/Abstract]) OR ("PD1"[All Fields] AND "suppressors"[Title/Abstract]) OR "anti-PD1"[Title/Abstract] OR "Nivolumab"[Title/Abstract] OR "Opdivo"[Title/Abstract] OR "ONO-4538"[Title/Abstract] OR "ONO4538"[Title/Abstract] OR "MDX-1106"[Title/Abstract] OR "MDX1106"[Title/Abstract] OR "BMS-936558"[Title/Abstract] OR "BMS936558"[Title/Abstract] OR "Pembrolizumab"[Title/Abstract] OR "SCH-900475"[Title/Abstract] OR "lambrolizumab"[Title/Abstract] OR "MK-3475"[Title/Abstract] OR "MK3475"[Title/Abstract] OR "Keytruda"[Title/Abstract] OR "Cemiplimab"[Title/Abstract] OR "REGN2810"[Title/Abstract] OR "Dostarlimab"[Title/Abstract] OR "Jemperli"[Title/Abstract] OR "dostarlimab-gxly"[Title/Abstract] OR "TSR-042"[Title/Abstract] OR "TSR042"[Title/Abstract] OR "Prolgolimab"[Title/Abstract] OR "Tislelizumab"[Title/Abstract] OR "BGB-A317"[Title/Abstract] OR "BGBA317"[Title/Abstract] OR "Retifanlimab"[Title/Abstract] OR "Durvalumab"[Title/Abstract] OR "MEDI4736"[Title/Abstract] OR "MEDI-4736"[Title/Abstract] OR "Imfinzi"[Title/Abstract] OR "Avelumab"[Title/Abstract] OR "MSB0010682"[Title/Abstract] OR "bavencio"[Title/Abstract] OR "MSB0010718C"[Title/Abstract] OR "MSB-0010718C"[Title/Abstract] OR "Atezolizumab"[Title/Abstract] OR "MPDL3280A"[Title/Abstract] OR "MPDL-3280A"[Title/Abstract] OR "Tecentriq"[Title/Abstract] OR "RG7446"[Title/Abstract] OR "RG-7446"[Title/Abstract] OR "Ipilimumab"[Title/Abstract] OR "Yervoy"[Title/Abstract] OR "MDX-010"[Title/Abstract] OR "MDX010"[Title/Abstract] OR "MDX-CTLA-4"[Title/Abstract] OR "MDXCTLA4"[Title/Abstract] OR "anti ctla 4 mab"[Title/Abstract] OR "Tremelimumab"[Title/Abstract] OR "CP-675"[Title/Abstract] OR "CP675"[Title/Abstract] OR "cp675 cpd"[Title/Abstract] OR "CP-675206"[Title/Abstract] OR "CP675206"[Title/Abstract] |
| #2 | "dement*"[Title/Abstract] OR "alzheimer*"[Title/Abstract] OR "lewy body"[Title/Abstract] OR "lewy bodies"[Title/Abstract] OR "lewy neurite"[Title/Abstract] OR "lewy neurites"[Title/Abstract] OR "amentia*"[Title/Abstract] OR "cognit*"[Title/Abstract] OR "learning and memory"[Title/Abstract] OR "glia*"[Title/Abstract] OR "astro*"[Title/Abstract] OR "tau"[Title/Abstract] OR "amyloid-beta"[Title/Abstract] OR "gliosis"[Title/Abstract] OR ("EORTC"[Title/Abstract] AND "QLQ"[Title/Abstract]) OR "european organization for research and treatment of cancer"[Title/Abstract] OR "european organisation for research and treatment of cancer"[Title/Abstract] OR "FACT-BRM"[Title/Abstract] OR "FACT-cog"[Title/Abstract] OR ("Neuropsychological"[Title/Abstract] AND "battery"[Title/Abstract]) OR "mini mental state exam*"[Title/Abstract] OR "MMSE"[Title/Abstract] OR "MOCA"[Title/Abstract] |
| #3 | #1 AND #2 |
| **Embase** | |
| #1 | ('pd-1':ab,ti OR 'pd1':ab,ti OR 'cd279':ab,ti OR 'pd-l1':ab,ti OR 'pdl1':ab,ti OR 'b7-h1':ab,ti OR 'b7h1':ab,ti OR 'cd274':ab,ti OR 'ctla-4':ab,ti OR 'ctla4':ab,ti OR 'cd152':ab,ti OR 'immune checkpoint':ab,ti OR 'immune checkpoints':ab,ti OR 'stimulatory checkpoint':ab,ti OR 'stimulatory checkpoints':ab,ti OR 'pd l1 inhibitor':ab,ti OR 'pd l1 inhibitors':ab,ti OR 'pd l1 blocker':ab,ti OR 'pd l1 blockers':ab,ti OR 'pd l1 suppressor':ab,ti OR 'pd-l1 suppressors':ab,ti OR 'pdl1 inhibitor':ab,ti OR 'pdl1 inhibitors':ab,ti OR 'pdl1 blocker':ab,ti OR 'pdl1 blockers':ab,ti OR 'pdl1 suppressor':ab,ti OR 'pdl1 suppressors':ab,ti OR 'anti-pdl1':ab,ti OR 'pd 1 inhibitor':ab,ti OR 'pd 1 inhibitors':ab,ti OR 'pd 1 blocker':ab,ti OR 'pd 1 blockers':ab,ti OR 'pd-1 suppressor':ab,ti OR 'pd-1 suppressors':ab,ti OR 'pd1 inhibitor':ab,ti OR 'pd1 inhibitors':ab,ti OR 'pd1 blocker':ab,ti OR 'pd1 blockers':ab,ti OR 'pd1 suppressor':ab,ti OR 'pd1 suppressors':ab,ti OR 'anti-pd1':ab,ti OR 'nivolumab':ab,ti OR 'opdivo':ab,ti OR 'ono-4538':ab,ti OR 'ono4538':ab,ti OR 'mdx-1106':ab,ti OR 'mdx1106':ab,ti OR 'bms-936558':ab,ti OR 'bms936558':ab,ti OR 'pembrolizumab':ab,ti OR 'sch-900475':ab,ti OR 'lambrolizumab':ab,ti OR 'mk-3475':ab,ti OR 'mk3475':ab,ti OR 'keytruda':ab,ti OR 'cemiplimab':ab,ti OR 'regn2810':ab,ti OR 'dostarlimab':ab,ti OR 'jemperli':ab,ti OR 'dostarlimab-gxly':ab,ti OR 'tsr-042':ab,ti OR 'tsr042':ab,ti OR 'prolgolimab':ab,ti OR 'tislelizumab':ab,ti OR 'bgb-a317':ab,ti OR 'bgba317':ab,ti OR 'retifanlimab':ab,ti OR 'durvalumab':ab,ti OR 'medi4736':ab,ti OR 'medi-4736':ab,ti OR 'imfinzi':ab,ti OR 'avelumab':ab,ti OR 'msb0010682':ab,ti OR 'bavencio':ab,ti OR 'msb0010718c':ab,ti OR 'msb-0010718c':ab,ti OR 'atezolizumab':ab,ti OR 'mpdl3280a':ab,ti OR 'mpdl-3280a':ab,ti OR 'tecentriq':ab,ti OR 'rg7446':ab,ti OR 'rg-7446':ab,ti OR 'ipilimumab':ab,ti OR 'yervoy':ab,ti OR 'mdx-010':ab,ti OR 'mdx010':ab,ti OR 'mdx-ctla-4':ab,ti OR 'mdxctla4':ab,ti OR 'anti ctla 4 mab':ab,ti OR 'tremelimumab':ab,ti OR 'cp-675':ab,ti OR 'cp675':ab,ti OR 'cp675 cpd':ab,ti OR 'cp-675206':ab,ti OR 'cp675206':ab,ti) AND [embase]/lim |
| #2 | ('dement*':ab,ti OR 'alzheimer*':ab,ti OR 'lewy body':ab,ti OR 'lewy bodies':ab,ti OR 'lewy neurite':ab,ti OR 'lewy neurites':ab,ti OR 'amentia*':ab,ti OR 'cognit*':ab,ti OR 'learning and memory':ab,ti OR 'glia*':ab,ti OR 'astro*':ab,ti OR 'tau':ab,ti OR 'amyloid-beta':ab,ti OR 'gliosis':ab,ti OR ('eortc':ab,ti AND 'qlq':ab,ti) OR 'european organization for research and treatment of cancer':ab,ti OR 'european organisation for research and treatment of cancer':ab,ti OR 'fact-brm':ab,ti OR 'fact-cog':ab,ti OR ('neuropsychological':ab,ti AND 'battery':ab,ti) OR 'mini-mental state exam*':ab,ti OR 'mmse':ab,ti OR 'moca':ab,ti) AND [embase]/lim |
| #3 | #1 AND #2 |
| **CENTRAL** | |
| #1 | ('pd-1':ab,ti OR 'pd1':ab,ti OR 'cd279':ab,ti OR 'pd-l1':ab,ti OR 'pdl1':ab,ti OR 'b7 h1':ab,ti OR 'b7h1':ab,ti OR 'cd274':ab,ti OR 'ctla-4':ab,ti OR 'ctla4':ab,ti OR 'cd152':ab,ti OR 'immune NEXT checkpoint':ab,ti OR 'immune NEXT checkpoints':ab,ti OR 'stimulatory NEXT checkpoint':ab,ti OR 'stimulatory NEXT checkpoints':ab,ti OR 'pd l1 inhibitor':ab,ti OR 'pd l1 inhibitors':ab,ti OR 'pd l1 blocker':ab,ti OR 'pd l1 blockers':ab,ti OR 'pd l1 suppressor':ab,ti OR 'pd-l1 suppressors':ab,ti OR 'pdl1 inhibitor':ab,ti OR 'pdl1 inhibitors':ab,ti OR 'pdl1 blocker':ab,ti OR 'pdl1 blockers':ab,ti OR 'pdl1 suppressor':ab,ti OR 'pdl1 suppressors':ab,ti OR 'anti-pdl1':ab,ti OR 'pd 1 inhibitor':ab,ti OR 'pd 1 inhibitors':ab,ti OR 'pd 1 blocker':ab,ti OR 'pd 1 blockers':ab,ti OR 'pd-1 suppressor':ab,ti OR 'pd-1 suppressors':ab,ti OR 'pd1 inhibitor':ab,ti OR 'pd1 inhibitors':ab,ti OR 'pd1 blocker':ab,ti OR 'pd1 blockers':ab,ti OR 'pd1 suppressor':ab,ti OR 'pd1 suppressors':ab,ti OR 'anti-pd1':ab,ti OR 'nivolumab':ab,ti OR 'opdivo':ab,ti OR 'ono-4538':ab,ti OR 'ono4538':ab,ti OR 'mdx-1106':ab,ti OR 'mdx1106':ab,ti OR 'bms-936558':ab,ti OR 'bms936558':ab,ti OR 'pembrolizumab':ab,ti OR 'sch-900475':ab,ti OR 'lambrolizumab':ab,ti OR 'mk-3475':ab,ti OR 'mk3475':ab,ti OR 'keytruda':ab,ti OR 'cemiplimab':ab,ti OR 'regn2810':ab,ti OR 'dostarlimab':ab,ti OR 'jemperli':ab,ti OR 'dostarlimab-gxly':ab,ti OR 'tsr-042':ab,ti OR 'tsr042':ab,ti OR 'prolgolimab':ab,ti OR 'tislelizumab':ab,ti OR 'bgb-a317':ab,ti OR 'bgba317':ab,ti OR 'retifanlimab':ab,ti OR 'durvalumab':ab,ti OR 'medi4736':ab,ti OR 'medi-4736':ab,ti OR 'imfinzi':ab,ti OR 'avelumab':ab,ti OR 'msb0010682':ab,ti OR 'bavencio':ab,ti OR 'msb0010718c':ab,ti OR 'msb-0010718c':ab,ti OR 'atezolizumab':ab,ti OR 'mpdl3280a':ab,ti OR 'mpdl-3280a':ab,ti OR 'tecentriq':ab,ti OR 'rg7446':ab,ti OR 'rg-7446':ab,ti OR 'ipilimumab':ab,ti OR 'yervoy':ab,ti OR 'mdx-010':ab,ti OR 'mdx010':ab,ti OR 'mdx-ctla-4':ab,ti OR 'mdxctla4':ab,ti OR 'anti ctla 4 mab':ab,ti OR 'tremelimumab':ab,ti OR 'cp-675':ab,ti OR 'cp675':ab,ti OR 'cp675 cpd':ab,ti OR 'cp-675206':ab,ti OR 'cp675206':ab,ti) |
| #2 | '('dement*':ab,ti OR 'alzheimer*':ab,ti OR 'lewy body':ab,ti OR 'lewy bodies':ab,ti OR 'lewy neurite':ab,ti OR 'lewy neurites':ab,ti OR 'amentia*':ab,ti OR 'cognit*':ab,ti OR 'learning and memory':ab,ti OR 'glia*':ab,ti OR 'astro*':ab,ti OR 'tau':ab,ti OR 'amyloid-beta':ab,ti OR 'gliosis':ab,ti OR ('eortc':ab,ti AND 'qlq':ab,ti) OR 'european organization for research and treatment of cancer':ab,ti OR 'european organisation for research and treatment of cancer':ab,ti OR 'fact-brm':ab,ti OR 'fact-cog':ab,ti OR ('neuropsychological':ab,ti AND 'battery':ab,ti) OR 'mini-mental state exam*':ab,ti OR 'mmse':ab,ti OR 'moca':ab,ti) |
| #3 | #1 AND #2 |
| **Web of Science** | |
| #1 | (TI=(PD-1 OR PD1 OR CD279 OR PD-L1 OR PDL1 OR B7-H1 OR B7H1 OR CD274 OR CTLA-4 OR CTLA4 OR CD152 OR immune NEXT/1 checkpoint OR immune NEXT/1 checkpoints OR stimulatory NEXT/1 checkpoint OR stimulatory NEXT/1 checkpoints OR PD-L1 inhibitor OR PD-L1 inhibitors OR PD-L1 blocker OR PD-L1 blockers OR PD-L1 suppressor OR PD-L1 suppressors OR PDL1 inhibitor OR PDL1 inhibitors OR PDL1 blocker OR PDL1 blockers OR PDL1 suppressor OR PDL1 suppressors OR anti-PDL1 OR PD-1 inhibitor OR PD-1 inhibitors OR PD-1 blocker OR PD-1 blockers OR PD-1 suppressor OR PD-1 suppressors OR PD1 inhibitor OR PD1 inhibitors OR PD1 blocker OR PD1 blockers OR PD1 blockers OR PDL suppressors OR anti-PD1 OR Nivolumab OR Opdivo OR ONO-4538 OR ONO4538 OR MDX-1106 OR MDX1106 OR BMS-936558 OR BMS936558 OR Pembrolizumab OR SCH-900475 OR SCH900475 OR lambrolizumab OR MK-3475 OR MK3475 OR Keytruda OR Cemiplimab OR REGN2810 OR Dostarlimab OR Jemperli OR dostarlimab-gxly OR TSR-042 OR TSR042 OR GSK4057190 OR Prolgolimab OR Tislelizumab OR BGB-A317 OR BGBA317 OR Retifanlimab OR Durvalumab OR MEDI4736 OR MEDI-4736 OR Imfinzi OR Avelumab OR MSB-0010682 OR MSB0010682 OR bavencio OR MSB0010718C OR MSB-0010718C OR Atezolizumab OR MPDL3280A OR MPDL-3280A OR Tecentriq OR RG7446 OR RG-7446 OR Ipilimumab OR Yervoy OR MDX-010 OR MDX010 OR MDX-CTLA-4 OR MDXCTLA4 OR Anti-CTLA-4 Mab OR Tremelimumab OR CP-675 OR CP675 OR CP675 cpd OR CP-675206 OR CP675206)) OR (AB=(PD-1 OR PD1 OR CD279 OR PD-L1 OR PDL1 OR B7-H1 OR B7H1 OR CD274 OR CTLA-4 OR CTLA4 OR CD152 OR immune NEXT/1 checkpoint OR immune NEXT/1 checkpoints OR stimulatory NEXT/1 checkpoint OR stimulatory NEXT/1 checkpoints OR PD-L1 inhibitor OR PD-L1 inhibitors OR PD-L1 blocker OR PD-L1 blockers OR PD-L1 suppressor OR PD-L1 suppressors OR PDL1 inhibitor OR PDL1 inhibitors OR PDL1 blocker OR PDL1 blockers OR PDL1 suppressor OR PDL1 suppressors OR anti-PDL1 OR PD-1 inhibitor OR PD-1 inhibitors OR PD-1 blocker OR PD-1 blockers OR PD-1 suppressor OR PD-1 suppressors OR PD1 inhibitor OR PD1 inhibitors OR PD1 blocker OR PD1 blockers OR PD1 blockers OR PDL suppressors OR anti-PD1 OR Nivolumab OR Opdivo OR ONO-4538 OR ONO4538 OR MDX-1106 OR MDX1106 OR BMS-936558 OR BMS936558 OR Pembrolizumab OR SCH-900475 OR SCH900475 OR lambrolizumab OR MK-3475 OR MK3475 OR Keytruda OR Cemiplimab OR REGN2810 OR Dostarlimab OR Jemperli OR dostarlimab-gxly OR TSR-042 OR TSR042 OR GSK4057190 OR Prolgolimab OR Tislelizumab OR BGB-A317 OR BGBA317 OR Retifanlimab OR Durvalumab OR MEDI4736 OR MEDI-4736 OR Imfinzi OR Avelumab OR MSB-0010682 OR MSB0010682 OR bavencio OR MSB0010718C OR MSB-0010718C OR Atezolizumab OR MPDL3280A OR MPDL-3280A OR Tecentriq OR RG7446 OR RG-7446 OR Ipilimumab OR Yervoy OR MDX-010 OR MDX010 OR MDX-CTLA-4 OR MDXCTLA4 OR Anti-CTLA-4 Mab OR Tremelimumab OR CP-675 OR CP675 OR CP675 cpd OR CP-675206 OR CP675206)) |
| #2 | (TI=(dement* OR alzheimer* OR "lewy body" OR "lewy bodies" OR "lewy neurite" OR "lewy neurites" OR amentia* OR cognit* OR "learning and memory" OR glia* OR astro* OR tau* OR "amyloid-beta" OR gliosis OR (eortc AND QLQ) OR "european organization for research and treatment of cancer" OR "european organisation for research and treatment of cancer" OR "fact-brm" OR "fact-cog" OR (neuropsychological AND battery) OR "mini-mental state exam*" OR MMSE OR MOCA)) OR (AB=(dement* OR alzheimer* OR "lewy body" OR "lewy bodies" OR "lewy neurite" OR "lewy neurites" OR amentia* OR cognit* OR "learning and memory" OR glia* OR astro* OR tau* OR "amyloid-beta" OR gliosis OR (eortc AND QLQ) OR "european organization for research and treatment of cancer" OR "european organisation for research and treatment of cancer" OR "fact-brm" OR "fact-cog" OR (neuropsychological AND battery) OR "mini-mental state exam*" OR MMSE OR MOCA)) |
| #3 | #1 AND #2 |

Supplementary Table 3. Characteristics of included preclinical studies with mouse models of Alzheimer’s disease

| First author, year | Subgroup | Mouse model  (n, sex, age) | | PD-1/PD-L1 modulation  (antibody, administration route,  dose, duration^1^) | Outcome | |
| --- | --- | --- | --- | --- | --- | --- |
|  |  | Control | Experimental |  | Cognitive behavioral change | Pathological change |
| Dvir-Szternfeld, 2022 | - | Trem2^+/+^5XFAD  + IgG  (10, M&F, 6-9 months) | Trem2^+/+^5XFAD  + anti PD-L1 antibody  (12, M&F, 6-9 months) | anti-PD-L1 antibody  (IP injection, 1.5mg/mouse, single injection, 1 month duration) | RAWM  NOR | N/A |
| Ben-Yehuda, 2021 | Ben-Yehuda 2021 | DM-Tau + IgG  (12, M&F, 8-13 months) | DM-Tau  + anti-PD-L1 antibody  (11, M&F, 8-13 months) | anti-PD-L1 antibody  (IP injection, 1.5mg/mouse, single injection, 1 month duration) | Y maze  NOR | N/A |
| Kummer, 2021 | - | APP/PS1  (7, NI, 9 months) | APP/PS1;PD-1 KO  (8, NI, 9 months) | Genetic deletion of PD-1 | MWM | Aβ plaque of hippocampus |
| Xing, 2021 | Xing 2021 B | APP/PS1 + IgG  (8, M, 8 months) | APP/PS1  + anti-PD-1 antibody  (8, M, 8 months) | anti-PD-1 antibody  (IP injection, 167ug/mouse, injected on Day 1 and 4, 1 month duration) | MWM | Aβ plaque of hippocampus |
|  | Xing 2021 C | APP/PS1 + IgG  (8, M, 8 months) | APP/PS1  + anti-PD-1 antibody  (8, M, 8 months) | anti-PD-1 antibody  (IP injection, 250ug/mouse, injected on Day 1 and 4, 1 month duration); | MWM | Aβ plaque of hippocampus |
| Zou, 2021 | Zou 2021 A | 5XFAD + IgG  (9, M, 9-10 months) | 5XFAD  + anti PD-1 antibody  (9, M, 9-10 months) | anti-PD-1 antibody  (ICV injection, 0.30 mg/mouse, single injection, 1 month duration) | MWM | Phosphorylated Tau (T231) |
|  | Zou 2021 B | WT  + Aβ insult  (8 M&F, 2-3months) | PD-1 KO  + Aβ insult  (8 M&F, 2-3 months) | Genetic deletion of PD-1 | N/A | Phosphorylated Tau (T231) |
|  | Zou 2021 C | WT  + Aβ insult  + IgG  (10, M&F, 2-3 months) | WT  + Aβ insult  + anti-PD-1 antibody  (10, M&F, 2-3 months) | anti-PD-1 antibody  (ICV injection, 0.3mg/mouse, single injection,12 days duration) | N/A | Phosphorylated Tau (T231) |
| Lin, 2020 | Lin 2020 | JNPL3tauopathy  (8, F, 13-14 months) | JNPLT3 tauopathy  + anti-PD-1 antibody  (10, F, 13-14 months) | anti-PD-L1 antibody  (IP injection, 10mg/kg, injected 12 times at 1 week interval, no duration time) | NOR  Y maze | N/A |
| Rosenzweig, 2019 | Rosenzweig  2019 A | 5XFAD + IgG  (9, NI 5.5 months) | 5XFAD  + anti-PD-1 antibody  (7, NI, 5.5months) | anti-PD-1 antibody  (IP injection, 0.5mg/mouse, injected on Day 1, 31 and 61, no duration time) | RAWM | N/A |
|  | Rosenzweig  2019 B | 5XFAD + IgG  (9, NI, 6.5 months) | 5XFAD  + anti-PD-1 antibody  (7, NI, 6.5 months) | anti-PD-1 antibody  (IP injection, 0.5mg/mouse, injected on Day 1, 31 and 61, 1 month duration) | RAWM | N/A |
|  | Rosenzweig  2019 C | 5XFAD + IgG  (15, NI, 7 months) | 5XFAD  + anti-PD-1 antibody  (14, NI, 7 months) | anti-PD-1 antibody  (i.p injection, 0.5 mg/mouse, single injection, 1 month duration) | RAWM | Aβ plaque of hippocampus  GFAP immunoreactivity |
|  | Rosenzweig  2019 D | 5XFAD + IgG  (15, NI, 7 months) | 5XFAD  + anti-PD-L1 antibody  (7, NI, 7 months) | anti-PD-L1 antibody  (IP injection, 0.5mg/mouse, single injection, 1 month duration) | RAWM | Aβ plaque of hippocampus  GFAP immunoreactivity |
|  | Rosenzweig  2019 E | DM-Tau + IgG  (11, M&F, 9 months) | DM-Tau  + anti-PD-1 antibody  (10, M&F, 9 months) | anti-PD-L1 antibody  (IP injection, 0.5mg/mouse, single injection, 1 month duration) | Y maze | Phosphorylated Tau (T231) |
|  | Rosenzweig  2019 F | DM-Tau + IgG  (11, M&F, 9 months) | DM-Tau  + anti-PD-L1 antibody  (7, M&F, 9 months) | anti-PD-L1 antibody  (IP injection, 0.5mg/mouse, single injection, 1 month duration) | Y maze | Phosphorylated Tau (T231) |
| Latta-Mahieu, | Latta-Mahieu 2017 A | APP/PS1 (ThyAPP/PS1M146L)  (9, M, 8 months) | APP/PS1 (ThyAPP/PS1M146L)  + anti-PD-1 antibody  (8, M, 8 months) | anti-PD-1 antibody  (IP injection, 10mg/kg, injection on Day 1, 4, 31, and 34, 1 month duration) | N/A | Aβ plaque of hippocampus |
|  | Latta-Mahieu 2017 B | APP/PS1 (ThyAPP/PS1A246E)  (17, NI, 11 months) | APP/PS1(ThyAPP/PS1A246E)  + anti-PD-1 antibody  (19, NI, 11 months) | anti-PD-1 antibody  (IP injection, 250g/mouse, injected on Day 1, 7, 29, and 32, 1 month duration) | N/A | Aβ plaque of hippocampus |
|  | Latta-Mahieu 2017 C | APP/PS1 (PD-APP)  (25, NI, 18 months) | APP/PS1(PD-APP)  + anti-PD-1 antibody  (25, NI, 18 months) | anti-PD-1 antibody  (IP injection, 8mg/kg, injected on Day 1, 4, 31, 34, 61, and 64, 20 days duration) | N/A | Aβ plaque of hippocampus |
| Baruch,2016 | Baruch 2016 A | 5XFAD + IgG  (6, M, 11 months) | 5XFAD  + anti PD-1 antibody  (9, M, 11 months) | anti-PD-1 antibody  (IP injection, 250ug/mouse, injected on Day 1 and 4, 1 month duration) | RAWM | N/A |
|  | Baruch 2016 B | 5XFAD + IgG  (9, M, 12 months) | 5XFAD  + anti PD-1 antibody  (4, M, 12 months) | anti-PD-1 antibody  (IP injection, 250ug/mouse, injected on Day 1 and 4, duration 2 months) | RAWM | Aβ plaque of hippocampus  GFAP immunoreactivity |
|  | Baruch 2016 C | 5XFAD + IgG  (9, M, 12 months) | 5XFAD  + anti PD-1 antibody  (6, M, 12 months) | anti-PD-1 antibody  (IP injection, 250ug/mouse, injected on Day 1, 4, 31 and 34, 1 month duration) | RAWM | Aβ plaque of hippocampus  GFAP immunoreactivity |
|  | Baruch 2016 D | APP/PS1  (4, M, 9 months) | APP/PS1  + anti-PD-1 antibody  (4, M, 9 months) | anti-PD-1 antibody  (IP injection, 250ug/mouse, injected on Day 1 and 4, 1 month duration) | N/A | Aβ plaque of hippocampus |
|  | Baruch 2016 E | APP/PS1  (4, F, 16 months) | APP/PS1  + anti-PD-1 antibody  (4, F, 16 months) | anti-PD-1 antibody  (IP injection, 250ug/mouse, injected on Day 1 and 4, 1 month duration) | N/A | Aβ plaque of hippocampus |

Abbreviations: Aβ, Amyloid beta; APP/PS1, Amyloid Precursor Protein-Presenilin 1; F, female; GFAP, Glial fibrillary acidic protein; IgG, Immunoglobulin G; IP, intraperitoneal; ICV, intracerebroventricular; KO, knock-out; M, male; MWM, Morris water maze; n, number; N/A, not applicable; NI, no-information; NOR, novel object recognition test; PD-1, programmed cell death protein 1; PD-L1, programmed cell death ligand 1; RAWM, Radial arm water maze; WT, wild-type; 5XFAD, 5-familiar Alzheimer’s disease;

^1^duration ; The time elapsed from the last injection to sacrifice or behavior experiment

Supplementary Table 4. Characteristics of included preclinical studies with wild-type mouse models

| First author, year | | Subgroup | Mouse model  (n, sex, age) | | | PD-1/PD-L1 modulation  (antibody, administration route, dose, duration) | Outcome |
| --- | --- | --- | --- | --- | --- | --- | --- |
|  |  |  | Control | Experimental | |  | Cognitive behavioral change |
| Zhao, 2023 | Zhao 2023 A | | WT  (11, M&F, 8-12 weeks) | | PD-1 KO  (11, M&F, 8-12 weeks) | Genetic deletion of PD-1 | MWM  NOR |
|  | Zhao 2023 B | | WT+IgG  (12, M&F, 8-16 weeks) | | WT  + anti-PD-1 antibody  (12, M&F, 8-16 weeks) | anti-PD-1 antibody  (ICV injection, 1ug/mouse, injected on  Day 1, 3 and 5 (+Day 7 and 9)^1^, 2 days duration) | MWM  NOR |
|  | Zhao 2023 C | | WT+IgG  (9, M&F, 8-16 weeks) | | WT  + anti-PD-L1 antibody  (9, M&F, 8-16 weeks) | anti-PD-L1 antibody  (ICV injection, 1ug/mouse,  injected 3 to 6 times at 2-day interval) | MWM  NOR |
| Kummer, 2021 | Kummer 2021 A | | WT  (12, NI, 9 months) | | PD-1 KO  (5, NI, 9 months) | Genetic deletion of PD-1 | MWM |
| Xing, 2021 | Xing 2021 A | | WT  (8, M, 8 months) | | WT  + anti-PD-1 antibody  (8, M, 8 months) | anti-PD-L1 antibody  (IP injection, 250ug/mouse, injected 2 times  at 3-day interval, 1 month duration); | MWM |
| Karl, 2019 | Karl 2019 | | WT  (14, M, 8-12 weeks) | | BH-71 KO  (14, M, 8-12 weeks) | Genetic deletion of BH-71 | MWM |
| Karl, 2017 | Karl 2017 A | | WT  (11, M&F, 8 weeks) | | BH-71 KO  (11, M&F, 8 weeks) | Genetic deletion of BH-71 | MWM |
|  | Karl 2017 B | | WT  (11, M&F, 12 months) | | BH-71 KO  (11, M&F, 12 months) | Genetic deletion of BH-71 | MWM |

Abbreviations: BH-71, B7 homolog 1; F, female; IgG, Immunoglobulin G; IP, intraperitoneal; ICV, intracerebroventricular; KO, knock-out; M, male; MWM, Morris water maze; n, number; NI, no-information; NOR, novel object recognition test; PD-1, programmed cell death protein 1; PD-L1, programmed cell death ligand 1; WT, wild-type;

^1^anti-PD-1 antibody was injected on day 1,3, and 5 for NOR test and additionally injected on Day 7 and 9 for MWM test

Supplementary Table 5. Characteristic of included clinical studies

| First author, year | Trial name | Phase | Follow-up period, weeks | Type of cancer | Line of therapy | Treatment group / Sample size | | | |
| --- | --- | --- | --- | --- | --- | --- | --- | --- | --- |
|  |  |  |  |  |  | Intervention | n | Comparison | n |
| Cescon, 2024 | KEYNOTE-355 | III | 15 | Triple-negative breast cancer | First-Line | Pembrolizumab+  Chemotherapy | 512 | Placebo+  Chemotherapy | 259 |
| Qin, 2024 | KEYNOTE-394 | III | 12 | Advanced hepatocellular carcinoma | Second-Line | Pembrolizumab | 298 | Placebo | 152 |
| Kim, 2023 | KEYNOTE-604 | III | 18 | Stage IV SCLC | First-Line | Pembrolizumab+  Chemotherapy | 208 | Placebo | 207 |
| Lorusso, 2023 | Study 309/  KEYNOTE-775 | III | 12 | Advanced Endometrial Cancer | NL | Pembrolizumab+  Lenvatinib | 370 | Doxorubicin or paclitaxel | 351 |
| Schmid, 2023 | KEYNOTE-119 | III | 6 | Metastatic triple-negative breast cancer | NL | Pembrolizumab | 289 | Capecitabine, eribulin, gemcitabine, or vinorelbine | 277 |
| Adenis, 2022 | KEYNOTE-181 | III | 9 | Advanced Esophageal Cancer | Second-Line | Pembrolizumab | 187 | Chemotherapy (paclitaxel, docetaxel, or irinotecan) | 188 |
| Oaknin, 2022 | EMPOWER CERVICAL-1 | III | 6 | Recurrent or metastatic cervical carcinoma | NL | Cemiplimab | 215 | Chemotherapy (pemetrexed, topotecan or irinotecan, gemcitabine, vinorelbine) | 181 |
| Rischin, 2022 | KEYNOTE-048 | III | 15 | Recurrent or metastatic head and neck squamous cell carcinoma | First-Line | Pembrolizumab alone | 280 | Cetuximab+  Chemotherapy | 262 |
|  |  |  |  |  |  | Pembrolizumab+chemotherapy | 255 | Cetuximab+  Chemotherapy | 244 |
| Zinzani, 2022 | KEYNOTE-204 | III | 24 | Relapsed or refractory Hodgkin lymphoma | NL | Pembrolizumab | 134 | Brentuximab vedotin | 138 |
| Andre, 2021 | KEYNOTE-177 | III | 18 | Colorectal cancer | First-Line | Pembrolizumab | 141 | Chemotherapy (mFOLFOX6 or FOLFIRI, with or without bevacizumab or cetuximab) | 131 |
| Harrington, 2021 | KEYNOTE-040 | III | 15 | Recurrent or metastatic head-and-neck squamous cell carcinoma | NL | Pembrolizumab | 231 | Chemotherapy (methotrexate, docetaxel, cetuximab) | 215 |
| Ryoo, 2021 | KEYNOTE-240 | II | 12 | Advanced Hepatocellular Carcinoma | Second-Line | Pembrolizumab | 271 | Placebo | 127 |
| Van Cutsem, 2021 (62) | KEYNOTE-062 | III | 18 | Advanced gastric/gastroesophageal junction adenocarcinoma | First-Line | Pembrolizumab | 239 | Placebo+  chemotherapy | 234 |
| Van Cutsem, 2021 (61) | KEYNOTE-061 | III | 12 | Advanced gastric/gastroesophageal junction cancer | Second-Line | Pembrolizumab | 173 | Paclitaxel | 170 |
| Garassino, 2020 | KEYNOTE-189 | III | 12 & 21 | Stage IV non-squamous NSCLC | First-Line | Pembrolizumab+  Chemotherapy (pemetrexed+  cisplatin or carboplatin) | 359 | Placebo+  Chemotherapy (pemetrexed+  cisplatin or carboplatin) | 180 |
| Mazieres, 2020 | KEYNOTE-407 | III | 9 & 18 | Stage IV squamous NSCLC | First-Line | Pembrolizumab+  Carboplatin-paclitaxel/nab-paclitaxel | 254 | Placebo+  Carboplatin-paclitaxel/nab-paclitaxel | 264 |
| Barlesi, 2019 | KEYNOTE-010 | II/III | 12 | NSCLC | NL | Pembrolizumab | 312 | Docetaxel | 266 |
| Vaughn, 2018 | KEYNOTE-045 | III | 15 | Advanced Urothelial Cancer | NL | Pembrolizumab | 260 | Chemotherapy (Paclitaxel, docetaxel, vinflunine) | 242 |
| Brahmer, 2017 | KEYNOTE-024 | III | 15 | Stage IV NSCLC | First-Line | Pembrolizumab | 145 | Chemotherapy | 137 |
| Harrington, 2017 | CheckMate 141 | III | 9 & 15 | Recurrent or metastatic head-and-neck squamous cell carcinoma | NL | Nivolumab | 93 | Chemotherapy (methotrexate, docetaxel, cetuximab) | 36 |
| Schadendorf, 2016 | KEYNOTE-002 | II | 12 | Unresectable stage III or IV melanoma | NL | Pembrolizumab | 169 | Chemotherapy | 170 |

Abbreviations: NL, not limited; NSCLC, non-small cell lung cancer; QLQ-C30, Quality of Life Questionnaire-Core-30; SCLC, small cell lung cancer

Supplementary table 6. Baseline characteristics of the study population in clinical studies

| First author, year | Age, median (range) | | Male, % | | ECOG PS, % | | Brian metastasis, n (%) | | | Dropout rate, % |
| --- | --- | --- | --- | --- | --- | --- | --- | --- | --- | --- |
|  | Intervention | Comparison | Intervention | Comparison | Intervention | Comparison | Intervention | | Comparison |  |
| Cescon, 2024 | 53.0 (44.0-63.0) | 53 (43.0-63.0) | 0 | 0 | 0: 59.0  1:41.0 | 0: 33.0  1: 38.0 | 17 (3.0) | 9 (3.0) | | 22 |
| Qin, 2024 | 54.0 (22.0-82.0) | 54.0 (22.0-78.0) | 85.7 | 82.4 | 0: 41.3 1: 58.7 | 0: 39.2 1: 60.8 | NI | | | 33 |
| Kim, 2023 | 64.0 (24.0-81.0) | 65.0 (37.0-83.0) | 66.7 | 63.1 | 0: 26.3 1: 73.7 | 0: 24.9 1: 75.1 | 33 (14.5) | 22 (9.8) | | 26 |
| Lorusso, 2023 | 64.0 (30.0-82.0) | 65.0 (35.0-86.0) | 0 | 0 | 0: 59.9 1: 39.9 | 0: 57.9 1: 42.1 | NI | | | 26 |
| Schmid, 2023 | 50.0 (43.0-59.0) | 53.0 (44.0-61.0) | 0 | 1 | 0: 54.0 1: 45.0 >=2: <1 | 0: 51.0 1: 49.0 >=2: 0 | 20 (6.0) | 22 (7.0) | | 13 |
| Adenis, 2022 | 63.0 (23.0-84.0) | 62.0 (24.0-84.0) | 86.9 | 86.3 | 0: 40.1 1: 59.6 2: 0.3 | 0: 36.9 1: 62.7 2: 0.3 | NI | | | 27 |
| Oaknin, 2022 | 51.0 (22.0-81.0) | 50.0 (24.0-87.0) | 0 | 0 | 0: 46.7 1: 53.3 | 0: 46.4 1: 53.6 | NI | | | 1 |
| Rischin, 2022^1^ | ^a^62.0 (56.0-68.0) | 61.0 (55.0-68.0) | 83.0 | 87.0 | 0: 39.0 1: 61.0 | 0: 40.0 1: 60.0 | NI | | | 19 |
|  | ^b^61.0 (55.0-68.0) | 61.0 (55.0-68.0) | 80.0 | 87.0 | 0: 39.0 1: 61.0 | 0: 39.0 1: 61.0 | NI | | | 15 |
| Zinzani, 2022 | 36.0 (28.0-53.0) | 35.0 (28.0-50.0) | 56.0 | 59.0 | 0: 57.0 1: 42.0 2: 1.0 | 0: 65.0 1: 35.0 2: 0 | NI | | | 37 |
| Andre, 2021^1^ | 63.0 (52.0-73.0) | 63.0 (50.0-72.0) | 46.1 | 52.1 | 0: 49.0 | 0: 55.0 | NI | | | 32 |
| Harrington, 2021^1^ | 60.0 (19.0-85.0) | 60.0 (34.0-78.0) | 15.4 | 17.5 | 0: 29.5 1: 70.5 | 0: 27.6 1: 72.4 | NI | | | 55 |
| Ryoo, 2021 | 67.0 (18.0-91.0) | 65.0 (23.0-89.0) | 81.3 | 83 | 0: 58.3 1: 41.7 | 0: 52.6 1: 47.4 | NI | | | 29 |
| Van Cutsem, 2021 (62) | 61.0 (20.0-83.0) | 62.5 (23.0-87.0) | 70.3 | 71.6 | 1: 48.8 | 1: 54.0 | NI | | | 49 |
| Van Cutsem, 2021 (61) | 62.5 (54.0-70.0) | 60.0 (53.0-68.0) | 68.0 | 70.0 | 0: 43.0 1: 57.0 2: 0 | 0:46.0 1: 53.0 2: <1 | NI | | | 42 |
| Garassino, 2020 | 65.0 (34.0-84.0) | 63.5 (34.0-84.0) | 62.0 | 52.9 | 0: 45.4 1: 53.9 2: 0.2 | 0: 38.8 1: 60.7 2: 0 | 73 (17.8) | 35 (17.0) | | 37 |
| Mazieres, 2020 | 65.0 (29.0-87.0) | 65.0 (36.0-88.0) | 79.1 | 83.6 | 0: 26.3 1: 73.7 | 0: 32.0 1: 68.0 | NI | | | 32 |
| Barlesi, 2019 | 63.0 (56.0-69.0) | 62.0 (56.0-69.0) | 62.0 | 61.0 | 0: 33.0 1: 67.0 2: 1.0 | 0: 34.0 1: 65.0 2: <1 | 56 (16.0) | | 48 (14.0) | 34 |
| Vaughn, 2018 | 67.0 (29.0-88.0) | 65.0 (26.0-84.0) | 74.1 | 74.3 | 0: 44.1 1: 53.0 2: 0.7 | 0: 39.0 1: 58.1 2: 1.5 | NI | | | 45 |
| Brahmer, 2017 | 64.5 (33.0-90.0) | 66.0 (38.0-85.0) | 59.7 | 62.9 | 0: 35.1 1: 64.3 | 0: 35.1 1: 64.9 | 18 (11.7) | | 10 (6.6) | 29 |
| Harrington, 2017^1^ | 61.0 (29.0-78.0) | 58.0 (39.0-74.0) | 82.1 | 85.1 | 0: 25.8 1: 74.2 | 0: 30.6 1: 69.4 | Excluded | | | 56 |
| Schadendorf, 2016 | 62.0 (15.0-87.0) | 63.0 (27.0-87.0) | 58.0 | 64.0 | 0: 54.0 1: 44.0 | 0: 55.0 1: 45.0 | Excluded | | | 32 |

Abbreviations: ECOG PS, The Eastern Cooperative Oncology Group performance status; NI, no information

^1^Baseline characteristics observed in patients who underwent health-related quality of life measurements

^2^Intervention group: pembrolizumab alone vs comparison group: cetuximab and chemotherapy

^3^Intervention group: pembrolizumab and cytotoxic chemotherapy vs comparison group: cetuximab and chemotherapy

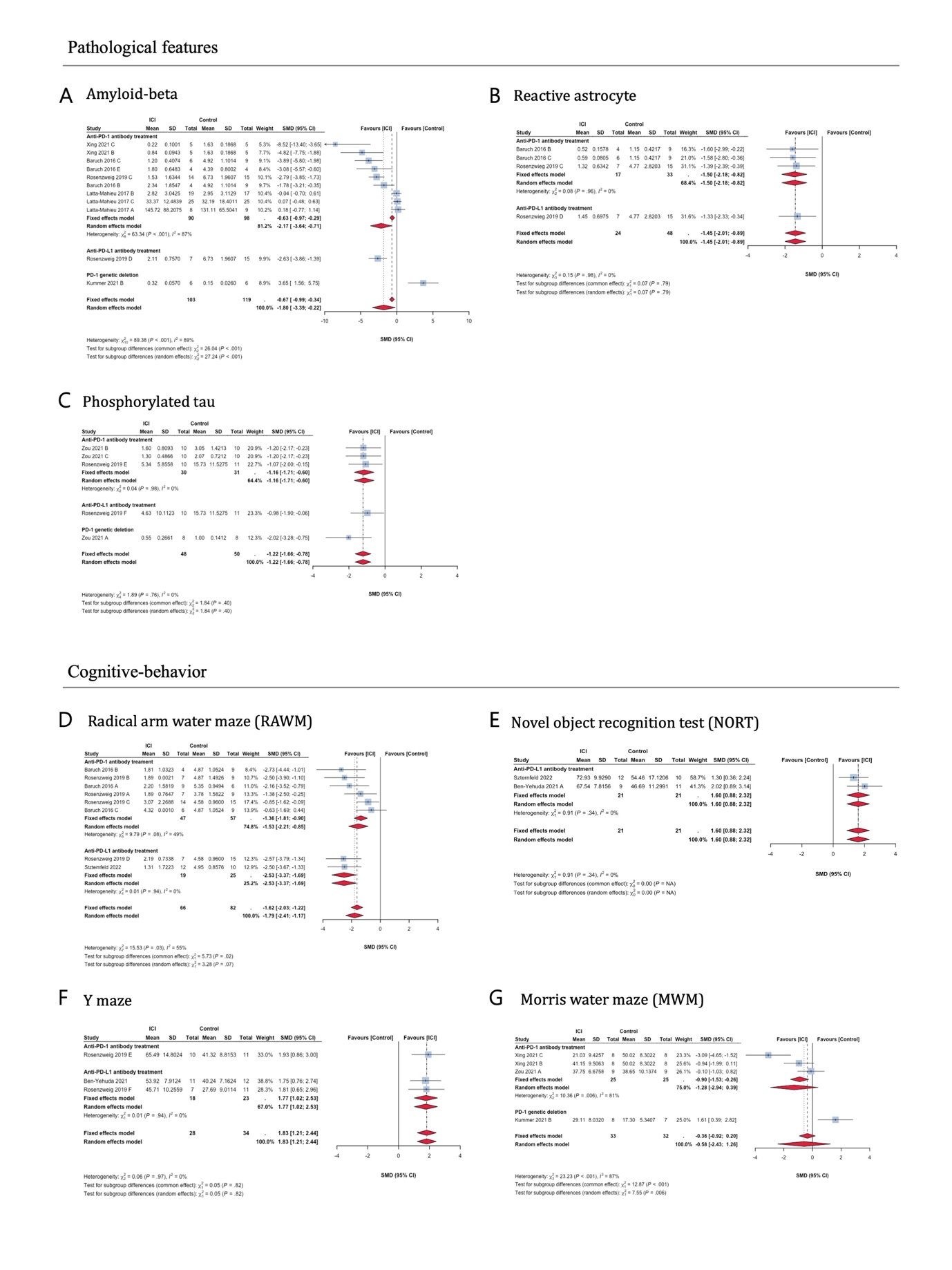

Supplementary figure 1. Detailed forest plot of pathological and cognitive-behavioral change in Alzheimer’s disease mouse models blocked by PD-1/PD-L1 signaling. (A) Amyloid-beta, (B) Reactive astrocyte, (C) Phosphorylated tau, (D) Radical arm water maze, (E) Novel object recognition test, (F) Y maze, (G) Morris water maze.

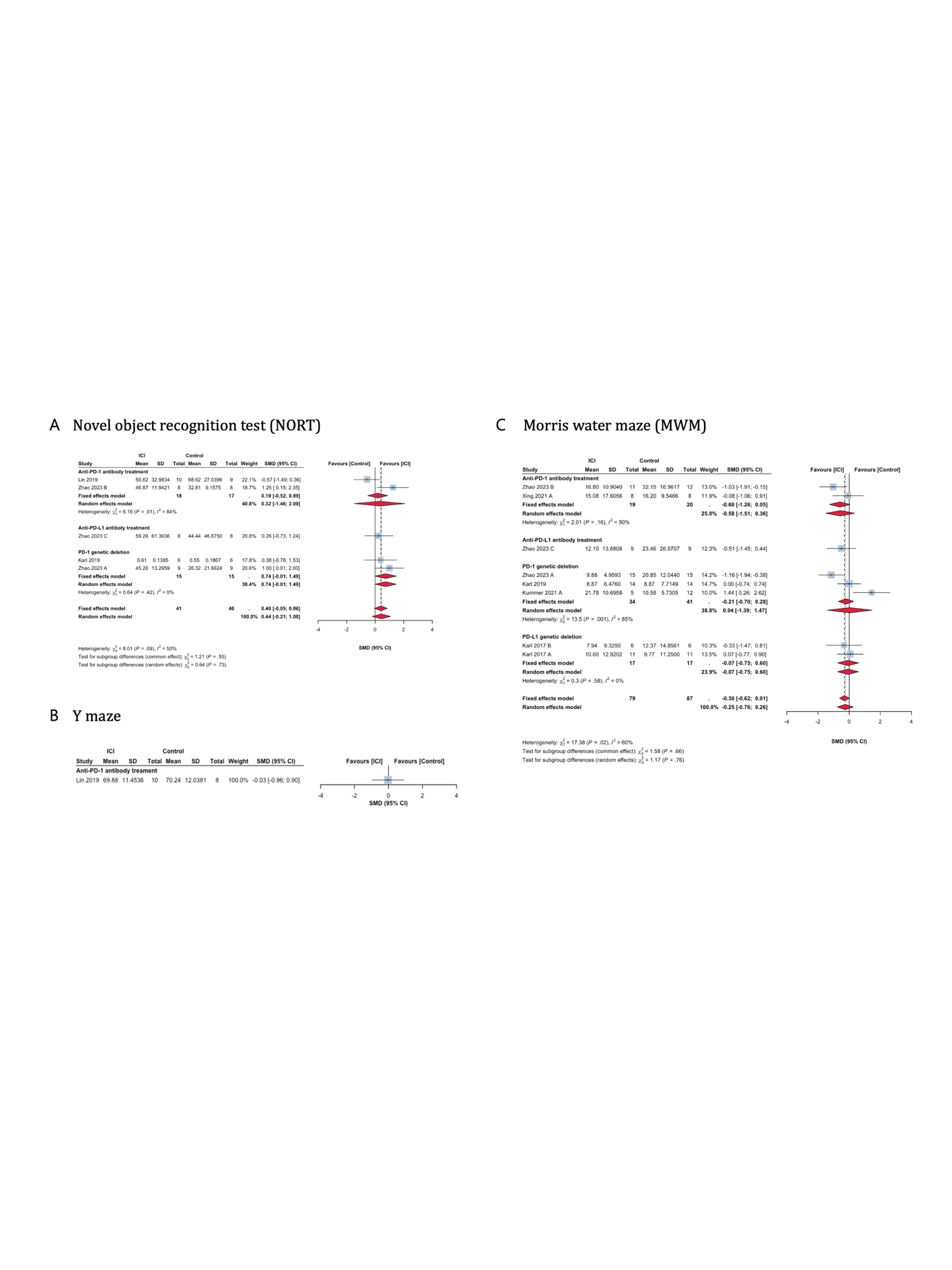

Supplementary figure 2. Detailed forest plot of cognitive behavioral change in wild-type mouse models blocked by PD-1/PD-L1 signaling. (A) Novel object recognition test, (B) Y maze, (C) Morris water maze.

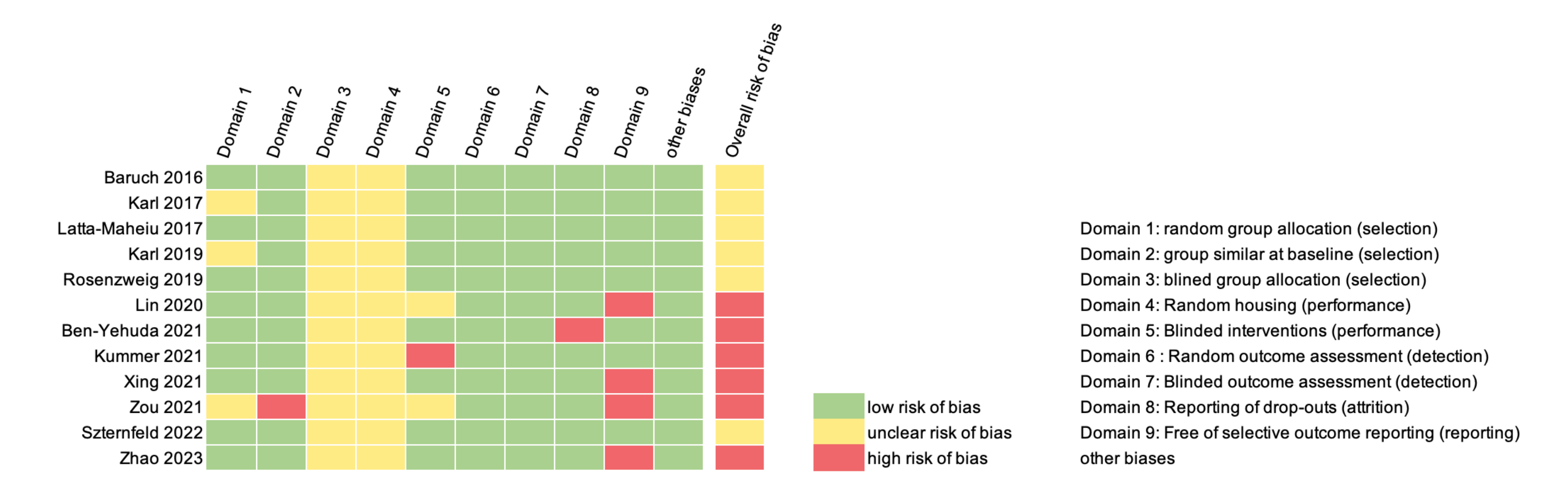

Supplementary figure 3. Quality assessment of the risk of bias according to the Systematic Review Centre for Laboratory Animal Experimentation criteria.

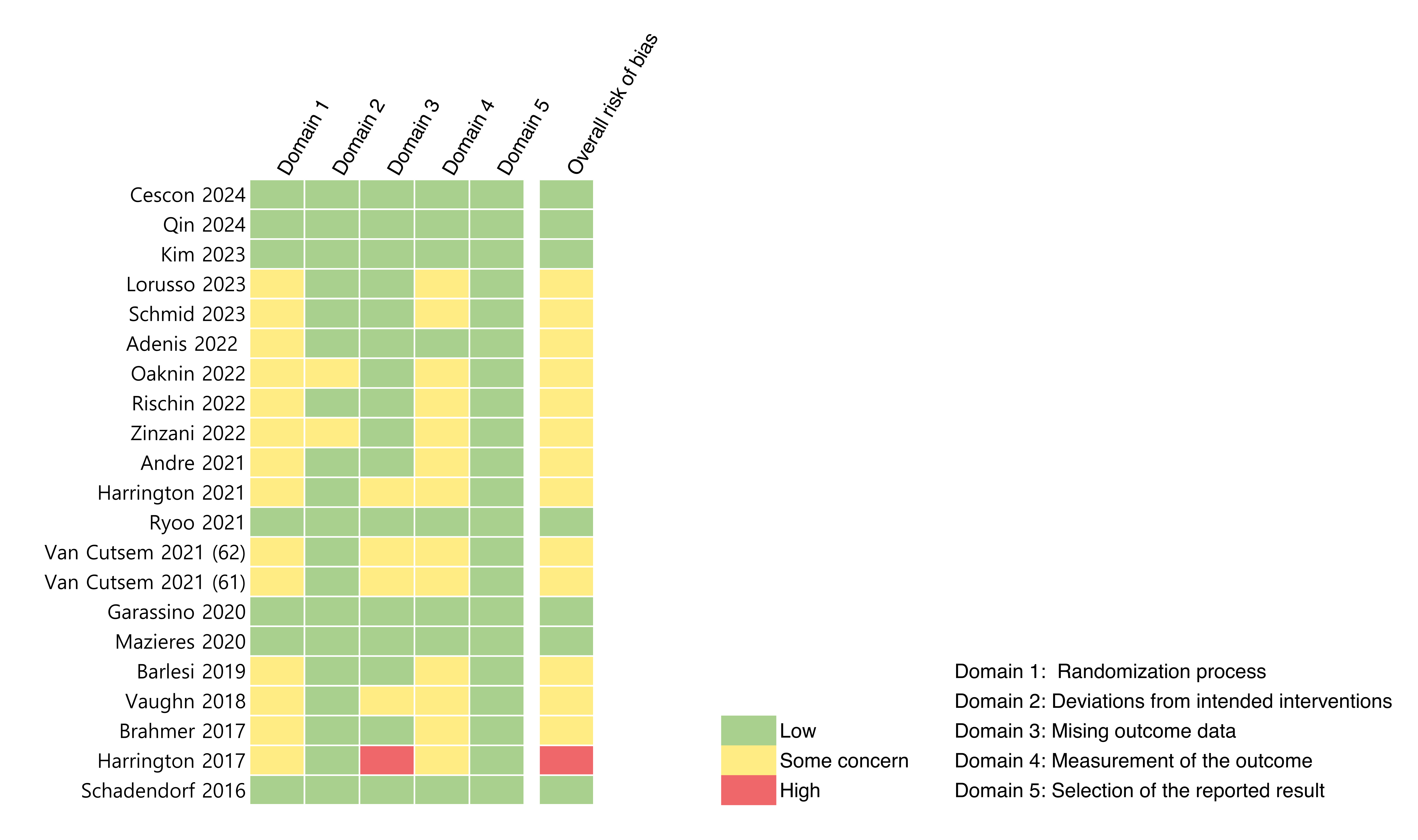

Supplementary figure 4. Quality assessment of the risk of bias according to the Risk of Bias 2 criteria.

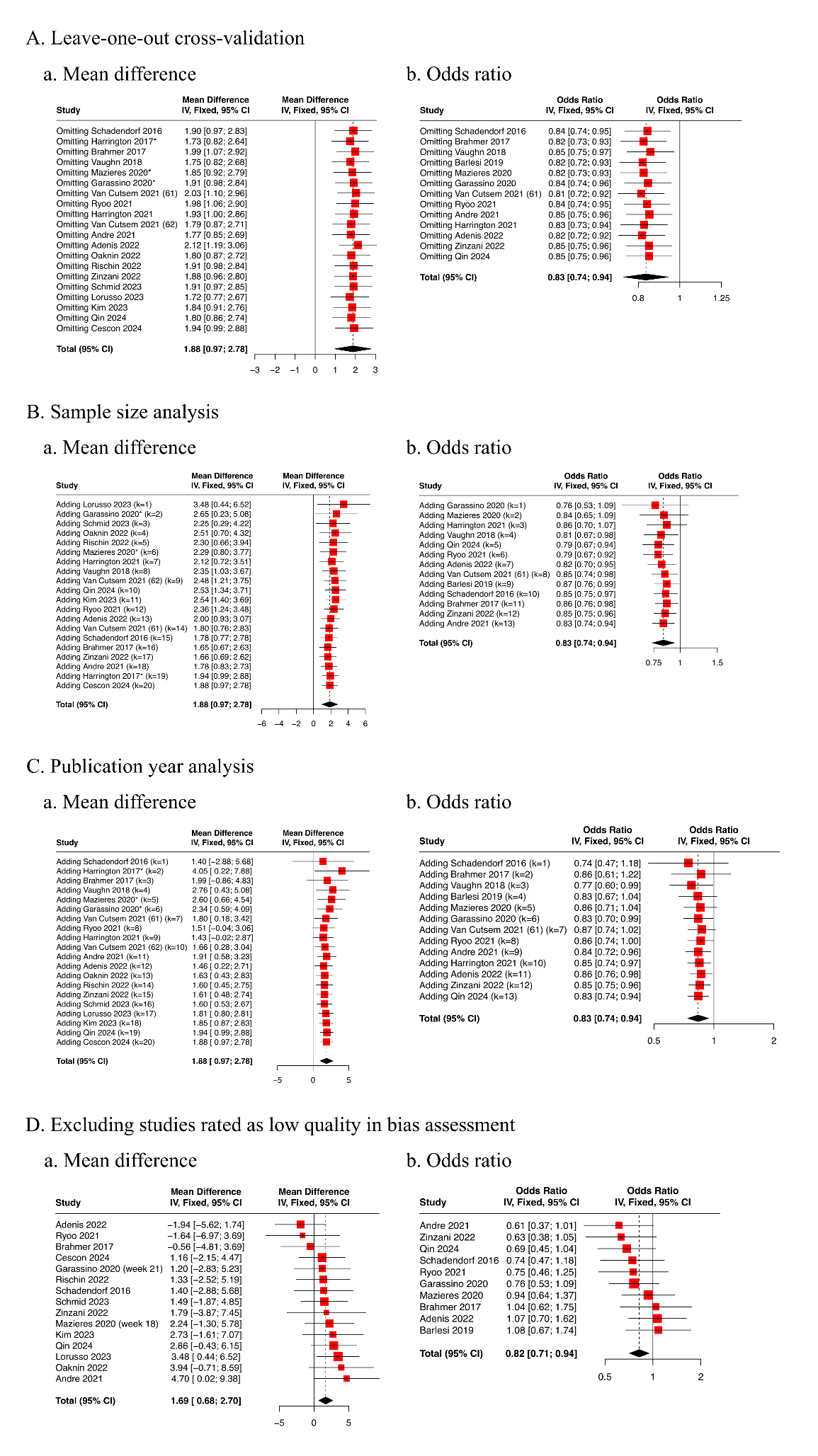

Supplementary figure 5. Sensitivity analysis of cognitive function change in clinical studies
